# Genetic variation at 11q23.1 confers colorectal cancer risk by dysregulation of colonic tuft cell transcriptional activator *POU2AF2*

**DOI:** 10.1101/2023.08.24.554659

**Authors:** V Rajasekaran, B. T Harris, R. T Osborn, C Smillie, K Donnelly, M Bacou, E Esiri-Bloom, L.Y Ooi, M Allan, M Walker, S Reid, A Meynert, G Grimes, J. P Blackmur, P. G Vaughan-Shaw, P. J Law, C Fernandez-Rozadilla, I. P Tomlinson, R Houlston, K. B Myant, F. V Din, M. G. Dunlop, S. M Farrington

**Author notes:** Contributed equally (V Rajasekaran, B. T Harris).

## Abstract

Common genetic variation at 11q23.1 is associated with colorectal cancer (CRC) risk, and exerts local (cis) expression quantitative trait locus (cis-eQTL) effects on *POU2AF2, COLCA1 and POU2AF3* genes. However, complex linkage disequilibrium and correlated expression at the 11q23.1 locus has thus far hindered elucidation of the mechanisms by which genetic variants impart CRC risk. Here, we establish that rs3087967 is the likely causal eQTL at this locus, co-localising with expression of *POU2AF2* and CRC risk. Furthermore, we show trans-eQTL effects on 21 distant target genes, which are highly enriched for Tuft cell markers. Analysis of available scRNAseq, ChIPseq and scATACseq data implicates POU2AF2 as the primary controller of the tuft cell specific trans-genes through POU2F3-correlated genetic regulation. Immunofluorescence demonstrates that the rs3087967 risk genotype (T) is associated with lower tuft cell abundance in human colonic epithelium. CRISPR-mediated deletion of the 11q23.1 risk locus in the mouse germline exacerbated the *Apc^Min/+^* mouse phenotype upon abrogation of *Pou2af2* expression specifically. Taken together, we implicate a key protective role of tuft cells in the large bowel and the importance of mis-regulation of *POU2AF2* as the prime tuft cell transcriptional activator at this locus.

## Introduction

Common, germline variation contributes up to 35% of overall colorectal cancer (CRC) risk^1–3^. To date, genome wide-association studies (GWAS) have identified 205 CRC risk-associated variants^4–6^, including rs3802842, a single nucleotide polymorphism (SNP) at 11q23.1^7^ (GRCh38 chr11:110,600,001-112,700,000). CRC-associated variation at 11q23.1 is corroborated by several studies; however, greater risk has been associated with alternative 11q23.1 variants, including rs11213801^8^, rs3087967^5,9^, rs7130173^10^ and rs10891245^10^. A lack of corroboration between these findings is likely due to high linkage disequilibrium (LD) at this region, making identification of the top-associated variant, an indicator of the target gene and potential dysregulatory mechanism, difficult.

Several studies also identify expression quantitative trait loci (eQTL) effects between CRC associated variation at 11q23.1 and downregulation of three local genes (cis-eQTL targets); *POU2AF2* (also known as *OCA-T1* or *C11orf53*), *COLCA1* and *POU2AF3* (also known as *OCA-T2* or *COLCA2*)^9–12^. Similarly, array expression analysis of healthy human colonic mucosa has identified numerous distant genes (trans-eQTL targets) to exhibit reduced expression with respect to variation at rs3087967 specifically, a SNP in the 3’ UTR of *POU2AF2*^13^. Our recent cell-specific mapping of 11q23.1 trans-eQTL target expression indicates these genes comprise a *POU2AF2*-correlated, tuft cell-specific transcriptional network, potentiating their status as putative markers of this cell-type and their expression to correlate with tuft cell abundance^14^. Correspondingly, POU2AF2 has recently been identified to interact with POU2F3, a master transcriptional regulator of tuft cells^15–18^, in a subtype of small cell lung cancer characterised by high *POU2F3* expression (SCLC-P)^19–21^. In addition, *Pou2af2* isoforms include variable domain composition in mice. The long *Pou2af2* isoform exclusively includes the putative POU2F interaction domain, recently mapped as the genetic locus associated with diversity of small intestinal tuft cell abundance across inbred mouse models^22^. While these studies potentiate tuft cell abundance to correlate with 11q23.1 eQTL target expression, genetic regulation of these genes, the presence and strength of this effect in the human colon, and it’s relevance to CRC risk are yet to be determined.

In this study, we interrogate genetic regulation of 11q23.1 eQTL targets in humans by analysing genome-wide molecular phenotypes at the bulk and single-cell level, assess cell abundance changes in the human colon, and experimentally delineate the function and tumourigenic potential of 11q23.1 cis-eQTL targets using genetically engineered mouse models.

## Results

### rs3087967 is the lead variant associated with 11q23.1 gene expression and CRC risk

To confirm 11q23.1 eQTL effects experimentally, we performed quantitative real-time PCR of *POU2AF2, COLCA1, POU2AF3* and the next closest gene, *POU2AF1,* across rs3802842 genotype in healthy colorectal mucosa, Supplementary Fig. S1. We find a significant cis-eQTL effect of rs3802842 for the expression of *POU2AF2* (p=0.0022), *COLCA1* (p=3.3e-09) and *POU2AF3* (p=1.5e-10), but not *POU2AF1* (p=0.787), thus validating the previously reported cis-eQTL effect of 11q23.1 variation^9–12^ and highlighting specificity to these genes.

Tagging SNPs are rarely found to be the causal variants in post-GWAS study, and as suggested previously, other 11q23.1 variants are more greatly associated with cis-eQTL target expression than rs3802842^9–12^. However, these studies used inferior expression quantification technologies that are smaller than current publicly available RNAseq datasets and genotyping arrays, which may miss more significant variants. To improve estimation of the relative effect of cis-eQTLs at 11q23.1, we performed linear regression of the expression of *POU2AF2, COLCA1* and *POU2AF3* and all detectable 11q23.1 variants in GTEx RNA-sequencing (RNAseq) and paired whole genome sequencing, focussing initially on the transverse colon (n=367), as GTEx sigmoid colon contains only muscularis, Fig 1 and Supplementary Material 1. We identified several variants to exhibit cis-eQTL effects that exceed the standard threshold of p<5e-8^6,23–25^, including previously reported variants rs11213801 (*POU2AF2* p=3.82e-15, *COLCA1* p=2.49e-16, *POU2AF3* p=2.88e-24), rs7130173 (*POU2AF2* p=1.09e-24, *COLCA1* p=1.72e-23, *POU2AF3* p=1.26e-37) and rs3087967 (*POU2AF2* p=1.27e-25, *COLCA1* p=2.00e-23, *POU2AF3* p=1.55e-39). In fact, based on absolute beta value, rs3087967 ranked second-highest amongst significant variants (p<5e-8) for *POU2AF2* expression and top for both *COLCA1* and *POU2AF3* expression, highlighting a particularly strong eQTL effect for this variant.

**Figure 1.**
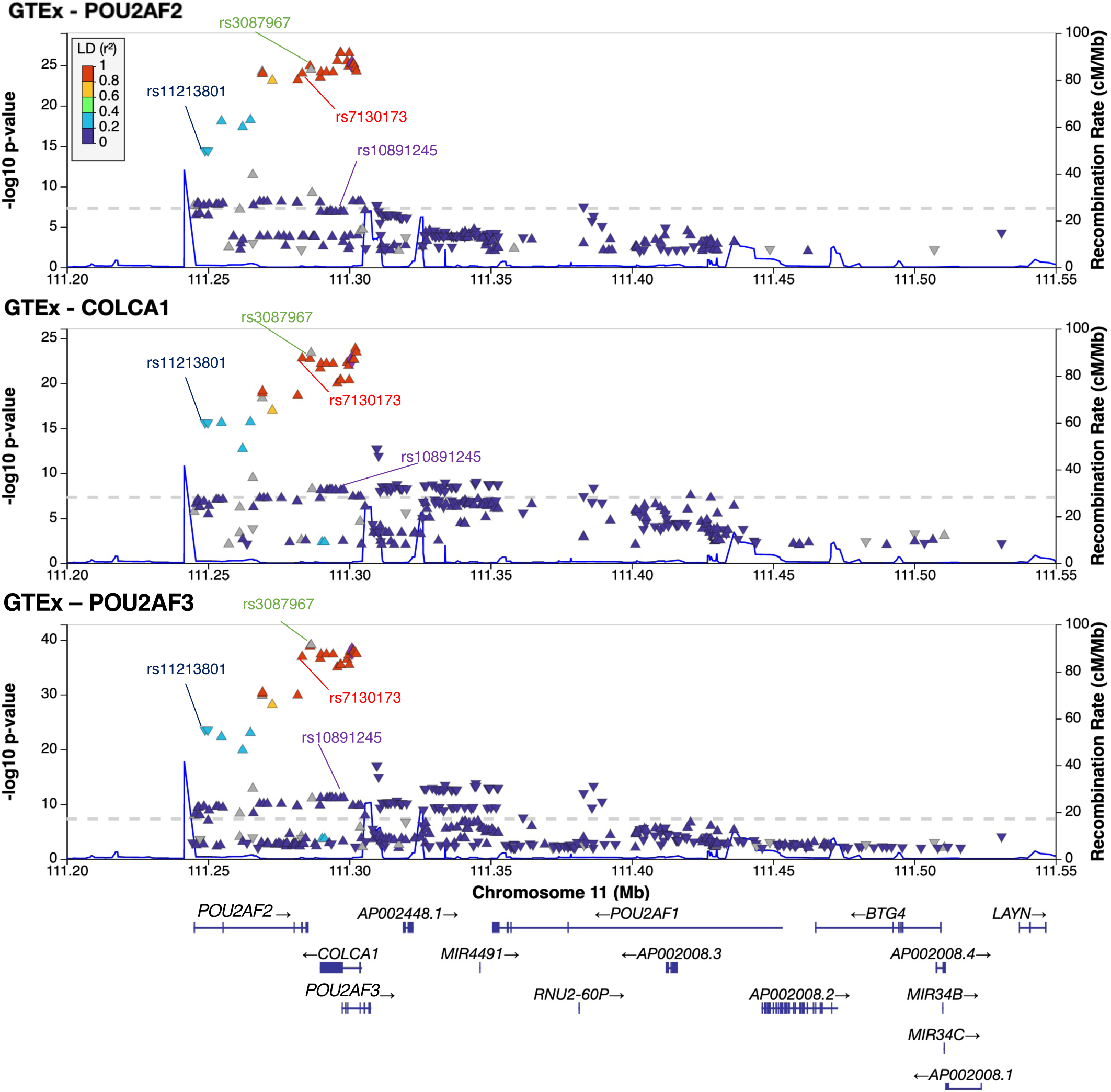
Several 11q23.1 variants exhibit significant associations with *POU2AF2, COLCA1* and *POU2AF3* expression in GTEx Transverse Colon RNAseq. locuzZoom^69^ plots of variant position and significance of association with *POU2AF2, COLCA1* and *POU2AF3* expression. Recombination rate (blue line) is derived from HapMap reference population. Variants are coloured by their LD with the tag variant (rs3802842), grey if no LD information available. Previously suggested top-associated variants^5,8,9,10^ are highlighted.

While this reinforces evidence for several eQTLs to be present at 11q23.1, it does not delineate this effect, or its association with CRC risk, to individual genes or variants. To investigate this, we tested the shared association of 11q23.1 variants with cis-eQTL target expression and CRC risk by colocalization analysis^26^, using summary statistics of our recent GWAS that identified rs3087967 as the top CRC risk variant at 11q23.1^6^, Supplementary Table S1. The expression of *POU2AF2, COLCA1* and *POU2AF3* was found to colocalize with CRC risk, with association at a single causal variant by far the most likely outcome in each case (Bayesian corrected posterior probability [PP]=0.91, 0.99 and 1 respectively). In addition, when investigating the 95% credible set of variants associated with both CRC risk and expression of each cis-eQTL target, rs3087967 was the only variant common to all three tests, with high significance for shared genetic association at a single causal variant in each case (PP= 0.75, 0.43, 0.99 respectively), Supplementary Table S2. To test whether there are independent variants to rs3087967 with a cis-eQTL effect, we performed conditional analysis, accounting for high LD at this region, finding no variant to be significantly, independently associated with expression of any 11q23.1 cis-eQTL target (conditional p-value < 5e-8), Supplementary Table S3. Together, this indicates 11q23.1 genetic variation is likely to represent a single cis-eQTL effect for *POU2AF2, COLCA1* and *POU2AF3* expression, which are all associated with CRC risk in bulk RNAseq. Importantly, this also highlights rs3087967 to be the most predictive variant for both eQTL and CRC risk effects.

### 11q23.1 variation does not exhibit transcript-specific eQTL effects

Recent work has not only highlighted a protein-protein interaction between POU2AF2 and POU2F3^19–21^, but also a dependence of murine small intestinal tuft cell abundance on expression of the POU2F interaction domain-encoding Pou2af2 transcript in mice^22^. Because of the homology between *POU2AF2*, *POU2AF3* and known POU2F interactor, *POU2AF1*^27–29^, we analysed the domain composition of proteins encoded by *POU2AF2* and *POU2AF3* transcripts in humans, Supplementary Fig S2. Of the two annotated *POU2AF2* transcripts expressed in the GTEx transverse colon, only ENST00000280325 includes the POU2F interaction domain, henceforth referred to as the ‘POU2F-ID transcript’. Meanwhile, two of the six *POU2AF3* transcripts expressed in GTEx transverse colon are POU2F-ID transcripts; ENST00000610738 and ENST00000638573. Interestingly, the only *POU2AF2* transcript associated with CRC risk in a recent transcript isoform wide association study was the POU2F-ID transcript^6^, potentially implicating this protein domain in governing 11q23.1 variation-associated CRC risk.

With scope to delineate eQTL effects to encoding of the POU2F-ID, we assessed transcript-specific eQTL effects of 11q23.1 variants in the colon. Interestingly, only the POU2F-ID transcript of *POU2AF2,* two non-POU2F-ID transcripts of *POU2AF3* and one POU2F-ID transcript of *POU2AF3* were significantly (p<5e-8) associated with rs3087967 genotype (p=1.05e-15, p=2.43e-15, p=1.33e-42 and p=3.41e-10 respectively). However, there is a clear trend for the reduced expression of the non-POU2F-ID *POU2AF2* transcript, indicating the lack of significance may be due to its lower overall expression. In contrast, rs3087967 and other 11q23.1 variants were significantly associated for both the POU2F-ID and non-POU2F-ID transcripts of *POU2AF3*, thus suggesting that CRC risk associated variation at 11q23.1 is not associated with perturbation of POU2F-ID transcripts of *POU2AF2* and *POU2AF3* specifically.

### Expression of twenty-one 11q23.1 trans-eQTL targets confer CRC risk

To better understand 11q23.1 trans-eQTL associations, we identified trans-eQTL targets of rs3087967 in GTEx transverse colon RNAseq, Supplementary Table S4, and two additional datasets; i) 223 in-house healthy colorectal mucosa samples by RNAseq (SOCCS^6^), Supplementary Table S5, and ii) 109 independent, healthy rectum mucosa samples by RNAseq (INTERMPHEN^5^), Supplementary Table S6. This replicated significant associations (FDR<0.05) with several targets previously identified^13^, including: *TRPM5; SH2D6; SH2D7; HTR3E; LRMP; GNG13; MATK; OGDHL; BMX* and *PLCG2*. Notably, we also identify *POU2F3* as a trans-eQTL target of rs3087967 (GTEx beta=0.58, FDR=1.39e-06), further potentiating altered tuft cell abundance in association with 11q23.1 genetic variation.

Replication of our eQTL analysis across all GTEx sites shows 11q23.1 variants exhibit significant association (FDR<0.05, beta>0.3) with each of *POU2AF2, COLCA1* and *POU2AF3* across numerous tissues, including; Esophagus Mucosa, Spleen, Nerve-Tibial and the Small Intestine-Terminal Ileum, Fig 2a. However, only in the transverse colon were >100 eQTLs identified for each of *POU2AF2*, *COLCA1* and *POU2AF3*. While an arbitrary threshold, that is subject to inherent variability in tissue-specific expression and dataset size, it suggests 11q23.1 cis-eQTL effects are strongest in the colon. Surprisingly, 11q23.1 trans-eQTL effects were only identified in the colon in these datasets, Fig 2b. The correlated exclusivity of 11q23.1 trans-eQTL effects and tumourigenic risk in this tissue may therefore imply causal relevance of the expression of these genes in governing CRC risk.

**Figure 2.**
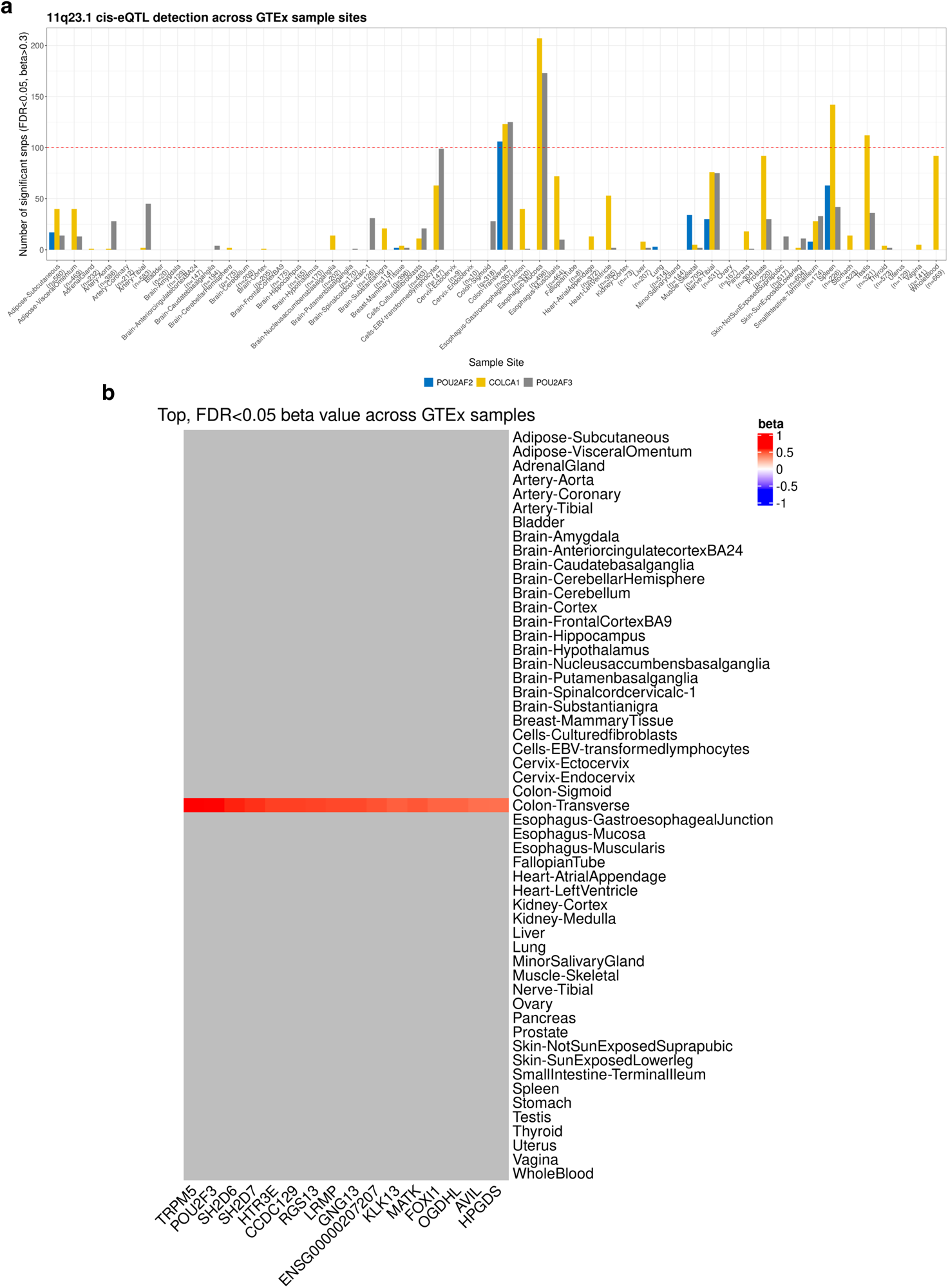
Pan-site 11q23.1 eQTL target analysis indicates risk-associated transcriptional dynamics are colon-specific. (a) Number of 11q23.1 variants significantly associated (FDR<0.05, beta>0.3) with each of *POU2AF2, COLCA1* and *POU2AF3* expression across all GTEx tissues (number of samples shown on x-axis). (b) Heatmap of the maximum, absolute beta value for significant (FDR<0.05) associations between 11q23.1 variants and rs3087967 trans-eQTL targets identified in the transverse colon (Table 1.4). Grey=No significant associations found.

To formally assess the causal potential of 11q23.1 trans-eQTL target expression in conferring CRC risk, we performed summarised mendelian randomisation (SMR), utilising genotype as an instrumental variable to estimate expression-associated risk^30^. After first meta-analysing 11q23.1 trans-eQTL associations across the two largest studies (GTEx transverse colon and SOCCS^6^), we limited our analysis to trans-eQTL targets with nominal significance in both studies (p<0.01, n=18,089). To protect against artificial inflation of SMR significance by constricting our analysis to 11q23.1, 11q23.1 trans-eQTLs were only used to perform SMR on trans-eQTL targets if the significance of their effect was greater than that of cis-eQTLs surrounding the trans-eQTL targets (n=1,474), Fig 3. Remarkably, the expression of twenty-one 11q23.1 trans-eQTL targets (trans-eQTL p<5e-8) were found to be associated with CRC risk at genome-wide significance (SMR p<8.4e-6), including *ACTG1P22, AVIL, AZGP1, B4GALNT4, CCDC129, CHAT, HCK, HPGDS, HTR3E, KLK13, LRMP, MATK, PIK3CG, PLCG2, POU2F3, PSTPIP2, RGS13, SH2D6, SH2D7, TAS1R3* and *TRPM5*, henceforth referred to as the ‘refined trans-eQTL targets’.

**Figure 3.**
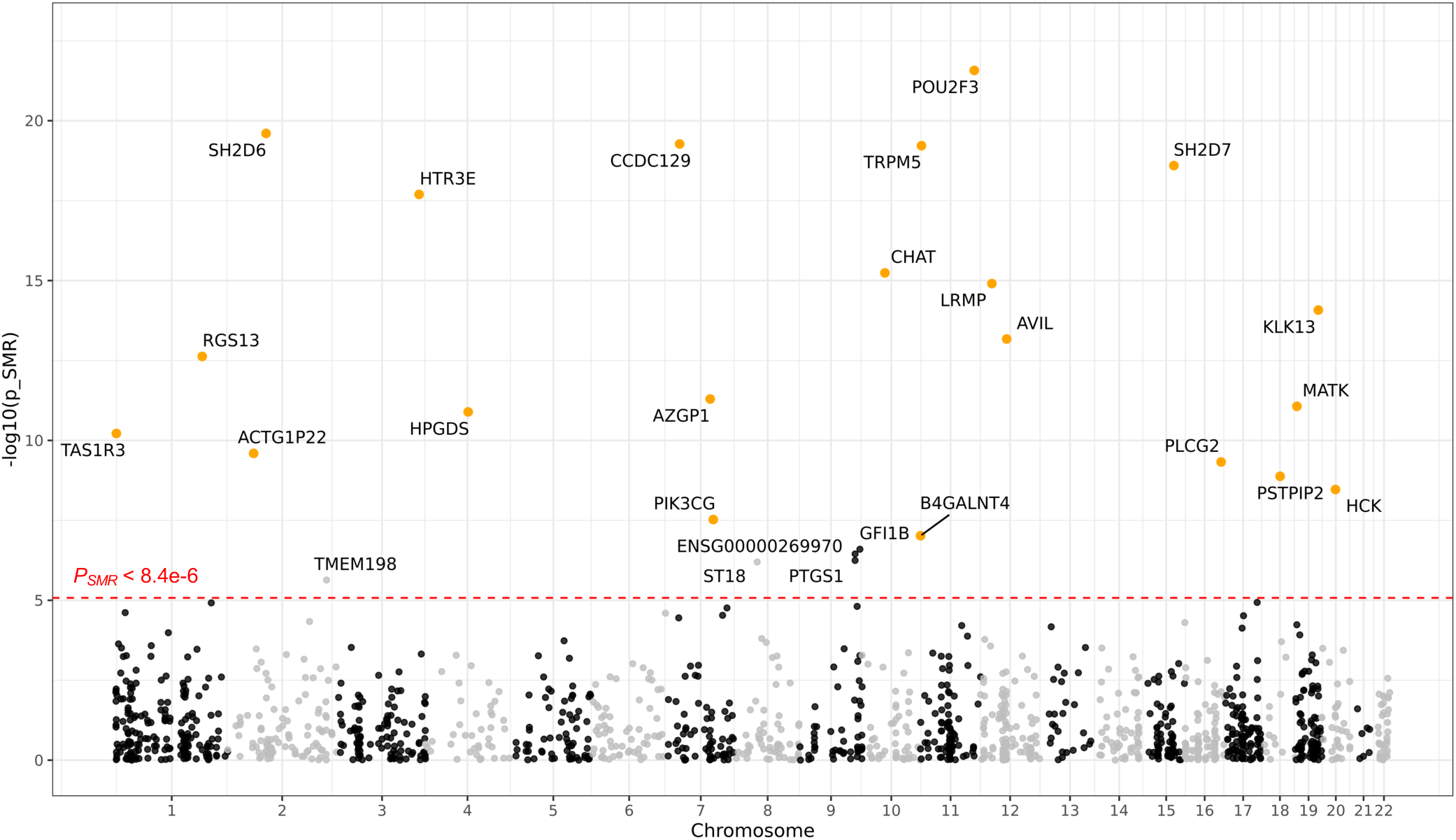
Summarised mendelian randomisation of 11q23.1 trans-eQTL targets identifies 21 novel genes with CRC risk association. SMR association between the expression of nominally significant 11q23.1 trans-eQTL targets (p<0.01) with greater 11q23.1 trans-eQTL effect than cis-eQTL effect (n=1,474) and CRC risk^6^. 11q23.1 trans-eQTL targets with genome-wide significance for CRC risk (pSMR<8.4e-6) are highlighted and orange for those with significant trans-eQTL effects (p < 5e-8).

Intriguingly, none of the refined trans-eQTL targets have been previously implicated as CRC risk associated by GWAS or transcriptome-wide association studies^5,6^. While other mechanisms may exist, it is possible that the expression of one or more 11q23.1 cis-eQTL targets directly or indirectly regulate the expression of these trans-eQTL targets. In this scenario, the expression of the trans-eQTL targets and their association with CRC risk would be dependent on both local eQTLs and the 11q23.1 trans-eQTLs. Such an example of epistasis would therefore hinder the identification of risk at the trans-eQTL target loci in case-control studies, that did not also take into account the 11q23.1 genotype. To address this, we performed an interaction, case-control analysis between cis-eQTLs within 1Mb of the refined trans-eQTL targets (bonferroni-corrected p-value<0.01) and rs3087967 genotype in a meta-analysis of Generation Scotland (n=14,205 [4,335 cases, 9,870 controls]) the Lothian Birth Cohort (n=2,550 [1,032 cases, 1518 controls]), and the National Study of Colorectal Cancer Genetics (n=13,801 [6,596 cases, 7,205 controls]). Interestingly, no variant within 1 Mb of these genes was found to be strongly associated with CRC risk based on interaction with rs3087967 genotype (minimum p=5.7e-4). We also performed a case-only interaction test in these studies, finding no evidence for cis-eQTLs surrounding the trans-eQTL targets, to be associated with CRC risk (minimum p=0.045). Together, this indicates epistatic effects of rs3087967 to not underpin the lack of CRC risk detected at these loci by GWAS. Instead, because cis-eQTL effects at these loci are of reduced significance compared with 11q23.1 trans-eQTL effects, this suggests the expression of these genes to be primarily regulated by 11q23.1 genotype, potentiating 11q23.1 cis-eQTL targets to be direct regulators of the refined trans-eQTL targets.

### POU2AF2 binding likely mediates tuft-cell specific accessibility of 11q23.1 trans-eQTL targets

While our RNAseq analyses reinforce the correlated expression of 11q23.1 cis- and trans-eQTL targets, it is not clear which, or what combination, of cis-eQTL targets are responsible for governing the trans effects. Our previous analysis of healthy human colonic epithelium single-cell RNAseq (scRNAseq) showed 11q23.1 trans-eQTL targets preferentially correlate with *POU2AF2* over *COLCA1* or *POU2AF3*^14^, but did not test the presently identified trans-eQTL target, *POU2F3,* or other CRC-risk-associated 11q23.1 trans-eQTL targets. Utilizing our previous analysis, we observed the mean expression of the refined trans-eQTL targets to be overwhelmingly greatest within tuft cells, including that of *POU2F3*, Supplementary Fig S3. In fact, 11 of the 17 refined trans-eQTL targets that passed gene filtration were identified as tuft cell markers in our previous analysis^14^ (Supplementary Table S7), and these preferentially correlate with one another and *POU2AF2*, over *COLCA1* or *POU2AF3,* in tuft cells only, Supplementary Fig S3. This therefore reinforces the specificity of the expression of the CRC associated 11q23.1 trans-eQTL targets, and the potential shared regulation with *POU2AF2*.

Given the recently identified interaction between POU2AF2 and POU2F3, and the transcriptional activator function of POU2AF2^19–21^, we sought to assess the co-binding of these proteins at refined 11q23.1 trans-eQTL targets as the potential mechanism by which their expression may be *POU2AF2*-regulated. To this end, we obtained publicly available chromatin immunoprecipitation sequencing (ChIPseq) of POU2AF2 and POU2F3 in SCLC-P cell lines (NCIH211 and NCIH526)^19^. While not colonic, or a tissue that preserves 11q23.1 trans-effects, lung epithelium is known to harbour tuft cells that express 11q23.1 trans-eQTL target genes^31^ and exhibit cis-eQTL associations between 11q23.1 genetic variation and *POU2AF2* expression (Fig 2a). POU2AF2 binding was identified at all refined trans-eQTL targets except for *ACTG1P22* and *CHAT,* while POU2F3 binding was identified at all except *ACTG1P22, CHAT* and *SH2D7* (Fig 4a). Binding at the refined trans-eQTL targets was significantly enriched (p<0.05) across all POU2AF2 ChIPseq replicates and the POU2F3 ChIPseq in NCIH211. Motif analysis of POU2AF2-bound 11q23.1 trans-eQTL targets also identified a striking enrichment of the POU2F3 binding motif (p=1e-127) at sixteen of the refined trans-eQTL targets, Fig 4b. Interrogation of sequence alignments at these regions highlights a specific, correlated pattern of POU2AF2 and POU2F3 binding (Supplementary Fig S4), complemented by active enhancer binding proteins: p300, MED1 (NCIH211 cell line only) and H3K27 acetylation (both NCIH211 and NCIH526 cell lines) at *AVIL, CCDC129, HCK, HTR3E, KLK13, LRMP, PIK3CG, PLCG2, POU2F3, PSTPIP2, RGS13, SH2D6* and *TAS1R3*. Together, this implicates POU2AF2 as a transcriptional activator of the majority of 11q23.1 trans-QTL gene targets by genomic binding in tandem with POU2F3, supporting this as the potential mechanism by which these genes are regulated. Interestingly, POU2AF2 and POU2F3 binding was also identified at *POU2AF2* and *POU2AF3* indicating POU2AF2 may participate in a positive feedback loop, and it’s expression may occur upstream of *POU2AF3*.

**Figure 4.**
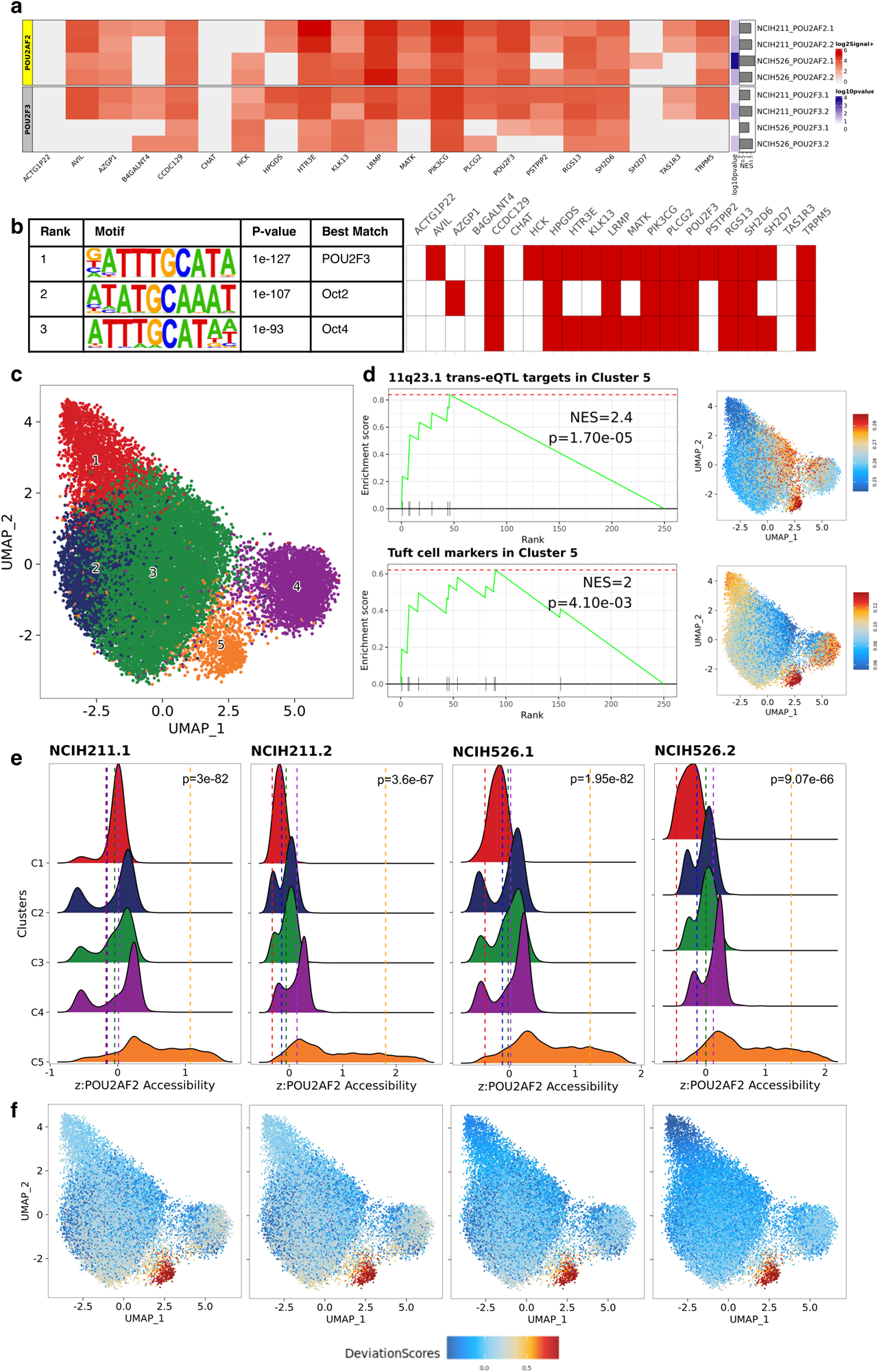
POU2AF2-enriched sequence accessibility is associated with colonic tuft cell chromatin accessibility landscape in the human colon. (a) Heatmap of signal values (log2+1) for POU2AF2 and POU2F3 binding at refined trans-eQTL targets across NCIH211 and NCIH526 SCLC-P cell lines. Gene set enrichment analysis of refined trans-eQTL targets in all bound sequences ranked by signal value also shown. (b) HOMER known motif enrichment results at POU2AF2 bound sequences surrounding refined 11q23.1 trans-eQTL targets (left). Heatmap denoting presence of core motif in sequences at refined trans-eQTL targets (right, red if present). (c) UMAP embedding of scATAC sequencing data from 21,620 healthy colonic epithelial cells^31^. Colour denotes cell-cluster. (d) Enrichment of refined 11q23.1 trans-eQTL targets and putative colonic tuft cell signature previously defined^14^ in cluster 5 accessibility marker genes and all cells. (e) Relative accessibility of POU2AF2-bound sequences for each antibody/cell line replicate across clusters and individual cells (f). Vertical line indicates the mean normalised accessibility for each cluster. P-values calculated by t-test of normalised enrichment scores in cluster 5 compared to all other clusters combined.

To test whether the potential genomic regulation of 11q23.1 trans-eQTL targets by POU2AF2 was also specific to tuft cells, we utilized single cell assay for transposase accessible sequencing (scATACseq) data of healthy human colonic epithelial cells (n=21,620)^32^. After dimensionality reduction and clustering, five distinct cell clusters were identified, Fig 4c. The genes with accessibility that demarcated cluster 5 were exclusively, significantly enriched for both the refined 11q23.1 trans-eQTL targets (NES=2.4, p=1.70e-05) and a putative tuft cell marker gene set (NES=2, p=4.10e-03), indicating the accessibility landscape of this cluster is likely characteristic of tuft cells. Accessibility of *PLCG2, LRMP, HCK, PSTPIP2, PIK3CG, B4GALNT4 TRPM5* and *HTR3E* demarcates cluster 5 (FDR<0.05), Supplementary Table S8, however several other refined trans-eQTL targets are highly cluster 5-specific, including; *AVIL, AZGP1, B4GALNT4, CHAT, HPGDS, KLK13, MATK* and *SH2D7,* Supplementary Fig S5. Analysis of healthy colonic scATACseq therefore supports the cell-specific expression of the majority of 11q23.1 trans-eQTL targets to be associated with corresponding cell-specific genomic regulation.

Master transcriptional regulators of differentiation, such as POU2F3, often function by modification of chromatin and transcriptional landscapes, yielding expression networks that direct specification of cell identity^33,34^. To agnostically test the potential for POU2AF2 binding to be associated with the tuft cell accessibility landscape, we analysed the relative enrichment of all POU2AF2-bound sequences across entire clusters and single cells, Fig 4e/f. We found a significant enrichment (maximum p=9.07e-66) of POU2AF2-bound regions in cluster 5 accessible regions (mean normalised accessibility range 1.07-1.8) compared with all other clusters (mean normalised accessibility range -0.49-0.15), highlighting a global enrichment of POU2AF2-bound sequence accessibility in cluster 5 specifically. Similarly, POU2F3-bound region accessibility was also significantly enriched in cluster 5 (mean normalised accessibility range 1.27-1.82 versus -0.44-0.13, maximum p=2.16e-63), Supplementary Fig S6. Together, this further reinforces evidence that POU2AF2 and POU2F3 coactivity regulates the colonic tuft cell genomic accessibility landscape and activates the tuft cell-specific expression of trans-eQTL targets, which is likely to be important in regulating tuft cell development.

#### CRC risk genotype at 11q23.1 is correlated with a reduced tuft cell compartment

As reduced expression of 11q23.1 trans-eQTL targets, including established (*POU2F3*) and putative tuft cell markers, are associated with 11q23.1 CRC risk (Supplementary Tables S4-6), we investigated tuft cell abundance in the healthy human colon across rs3087967 genotype. Because of the known rarity of tuft cells, we sought to minimise variation in cell abundance due to technical sampling effects, by maximising the proportion of epithelium on histological slides. To this end, we optimised a novel method of human intestinal sample collection by rolling stripped epithelium from colonic mucosa to produce ‘swiss rolls’, approximately six of which can be included on a single 25mm x 75mm histological slide. Each ‘swiss roll’ is comprised of a dramatic enrichment (p=4.42e-05) of epithelial/stromal cell density (mean=2001/mm^2^, median=1995/mm^2^, 95% confidence interval=1709-2558/mm^2^) compared to non-rolled tissue (mean=658/mm^2^, median=482/mm^2^, 95% confidence interval=311-1007/mm^2^), Supplementary Fig S7, Video S1 and S2. To assess tuft cell abundance across rs3087967 genotype, we performed dual immunofluorescence of additional tuft cell markers ChAT^35,36^ and COX1/PTGS1^36,37^ with POU2F3 in colonic swiss rolls, Fig 5. In accordance with risk associated expression dynamics, we identified a significant reduction in the relative abundance of tuft cells in healthy colonic mucosa of individuals that were homozygous for the CRC risk allele at rs3087967 (TT, n=4) for both POU2F3/ChAT (p=0.042) and POU2F3/PTGS1 (p=0.042) compared to samples that were homozygous non-risk (CC, n=7). This therefore confirms genetic variation at 11q23.1, associated with CRC risk, is also correlated with reduced tuft cell abundance in the human colon.

**Figure 5.**
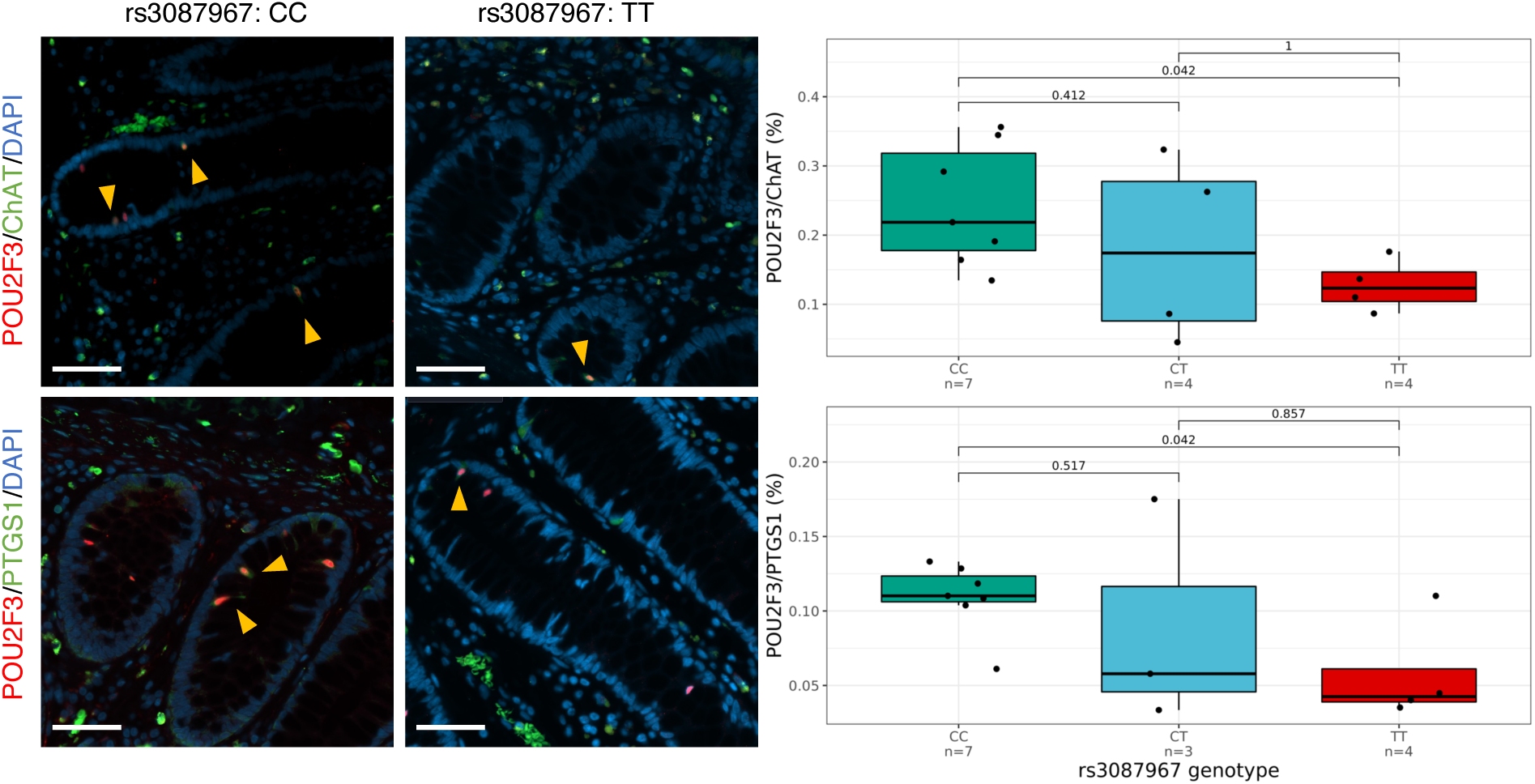
CRC risk genotype at 11q23.1 is associated with reduced colonic tuft cell abundance. Double immunofluorescent staining of tuft cell markers choline acetyltransferase (ChAT)/POU2F3 (top) and PTGS1/POU2F3 (bottom) in human colon epithelium across rs3087967 genotype, C=non-CRC risk allele, T=CRC risk-associated allele. P-values calculated by unpaired Wilcoxon test. Scale=50μm. Example positive cells identified with yellow arrowheads.

#### Murine 11q23.1 gene knockout models human transcriptional dynamics, cell abundance changes and yield diverse phenotypes

While the ChIPseq, scATACseq and scRNAseq analyses imply colonic tuft cell genomic accessibility and transcriptional landscapes are dependent on *POU2AF2* expression, we cannot exclude potential associations between the abundance of this cell type with *COLCA1* or *POU2AF3.* Unfortunately, all three genes are poorly expressed across colorectal cancer cell lines and highly correlated in human bulk expression (Supplementary Fig S1) making experimental delineation of tuft cell abundance and potential tumourigenesis in human tissue or models difficult. To this end, we reasoned genetic perturbation of individual or combinations of 11q23.1 homologs in mice would allow direct interrogation of gene function and associated phenotypes.

To validate mice as a model of human 11q23.1-associated transcriptional dynamics, we generated a knockout model of all cis-eQTL target homologs in C57/Bl6 mice via CRISPR-Cas9 (Fig 6a), hereafter referred to as *C11orfΔ*. We performed bulk RNAseq of proximal and distal colonic samples from *C11orfΔ^-/-^* (n=4) and wild-type mice (n=7), identifying significantly reduced expression of *Pou2af2, Pou2af3* and homologs of refined 11q23.1 trans-eQTL targets: *Sh2d7, Trpm5, Sh2d6, Pou2f3, Ccdc129, Rgs13* and *Avil,* in addition to non-refined trans-eQTL target homolog, *Alox5* (FDR<0.05, logFC<-1.5), Fig 6b. We also analysed the expression of individual transcripts across *C11orfΔ* genotype. Each of *Pou2af2* and *Pou2af3* encode only a single POU2F-ID transcript^22^ (Fig 6c), but all *Pou2af2* and *Pou2af3* transcripts that passed quality control (see Methods) were significantly reduced in *C11orf11^-/-^* mice (FDR<0.05), Fig 6d. The correlated downregulation of both cis- and trans-eQTL target homologs in this model highlights a striking similarity to the transcriptional dynamics associated with 11q23.1 genetic variation in humans, and while this does not offer the resolution to delineate this effect to encoding of the Pou2f interaction domain, it validates the use of mice to model this CRC risk locus.

**Figure 6.**
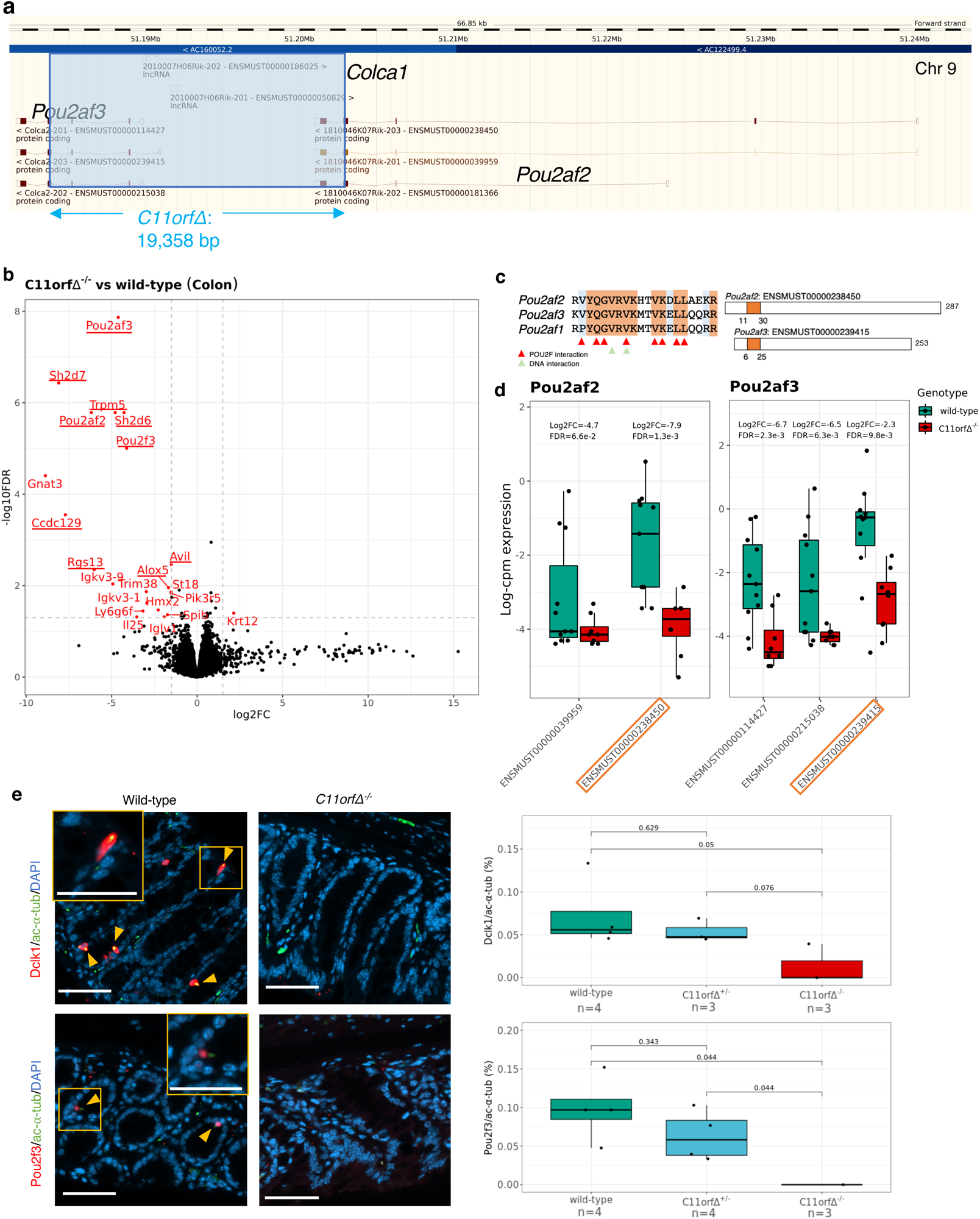
*C11orfΔ^-/-^* mice recapitulate 11q23.1 variation associated transcriptional dynamics and tuft cell abundance. (a) Generation of the *C11orfΔ* model, characterised by a 19,358 bp deletion across all three homologs of 11q23.1 cis-eQTLs. Schematic obtained from ensembl (https://www.ensembl.org/, accessed 21/09/22) (b) Volcano plot of the differentially expressed (DE) genes from *C11orfΔ^-/-^* (n=4) vs wild-type (n=7) colon RNAseq. Genes with absolute log_2_ fold-change > 1.5 and FDR<0.05 are highlighted in red and underlined if an 11q23.1 trans-eQTL target homolog. (c) Pou2f interaction domain homology between Pou2af1, Pou2af2 and Pou2af3 proteins (left) and transcripts encoding this domain (right and boxed to highlight). (d) Expression of detected *Pou2af2* and *Pou2af3* transcripts across *C11orfΔ^-/-^* and wild-type colon RNAseq. False Discovery Rate (FDR) calculated to account for the number of cis-eQTL transcripts being tested only. Orange highlight indicates transcripts encoding Pou2f interaction domain. (e) Immunofluorescent stains of Dclk1/acetylated-⍺-tubulin (ac-⍺-tub) and Pou2f3/ac-⍺-tub across *C11orfΔ^-/-^* genotype. Scale bars are 50μm. P-values calculated by unpaired Wilcoxon.

While *C11orf11^-/-^* mice did not harbour tumours in the colon or elsewhere, several phenotypes were observed. *C11orf11^-/-^* mice consistently weighed less than *C11orf11^+/-^* and wild-type littermates and showed reduced overall survival, indicative of reduced thriving that may be linked to impaired intestinal function, Supplementary Fig S8. In addition, male *C11orf11^-/-^* mice were infertile and while this is not likely to represent processes associated with CRC risk, it does indicate potential functions of 11q23.1 cis-eQTL targets in other tissues.

To assess the abundance of tuft cells in these mice, we performed immunofluorescence of both Dclk1 and Pou2f3 in tandem with acetylated alpha tubulin (ac-α-tub), Fig 6e. Analogous to the human colon, we observed a significant depletion in the relative abundance of Dclk1/ac-α-tub and Pou2f3/ac-α-tub cells in *C11orf11^-/-^* colon (n=3) compared with wild-type (n=4; p=0.05, p=0.044 respectively). This therefore also shows our genetically engineered mouse models recapitulate the human tuft cell abundance changes observed across 11q23.1 genotype.

#### Colonic tuft cell abundance is dependent on *Pou2af2,* not *Pou2af3* expression

To interrogate the contribution of individual 11q23.1 cis-eQTL target homologs to *C11orf11^-/-^* transcriptional dynamics and phenotypes, we generated an additional knockout mouse model of *Pou2af2* by CRISPR-Cas9, and obtained a *Pou2af3* model from the Canadian Mutant Mouse Repository^38^. The *Pou2af2* knockout model is comprised of an 11bp deletion of exon 4, resulting in a frameshift mutation that affects all *Pou2af2* transcripts, Fig 7a. Meanwhile the *Pou2af3* model is characterised by a 689bp deletion spanning the entire third exon, common to all *Pou2af3* transcripts, and results in a premature stop codon.

**Figure 7.**
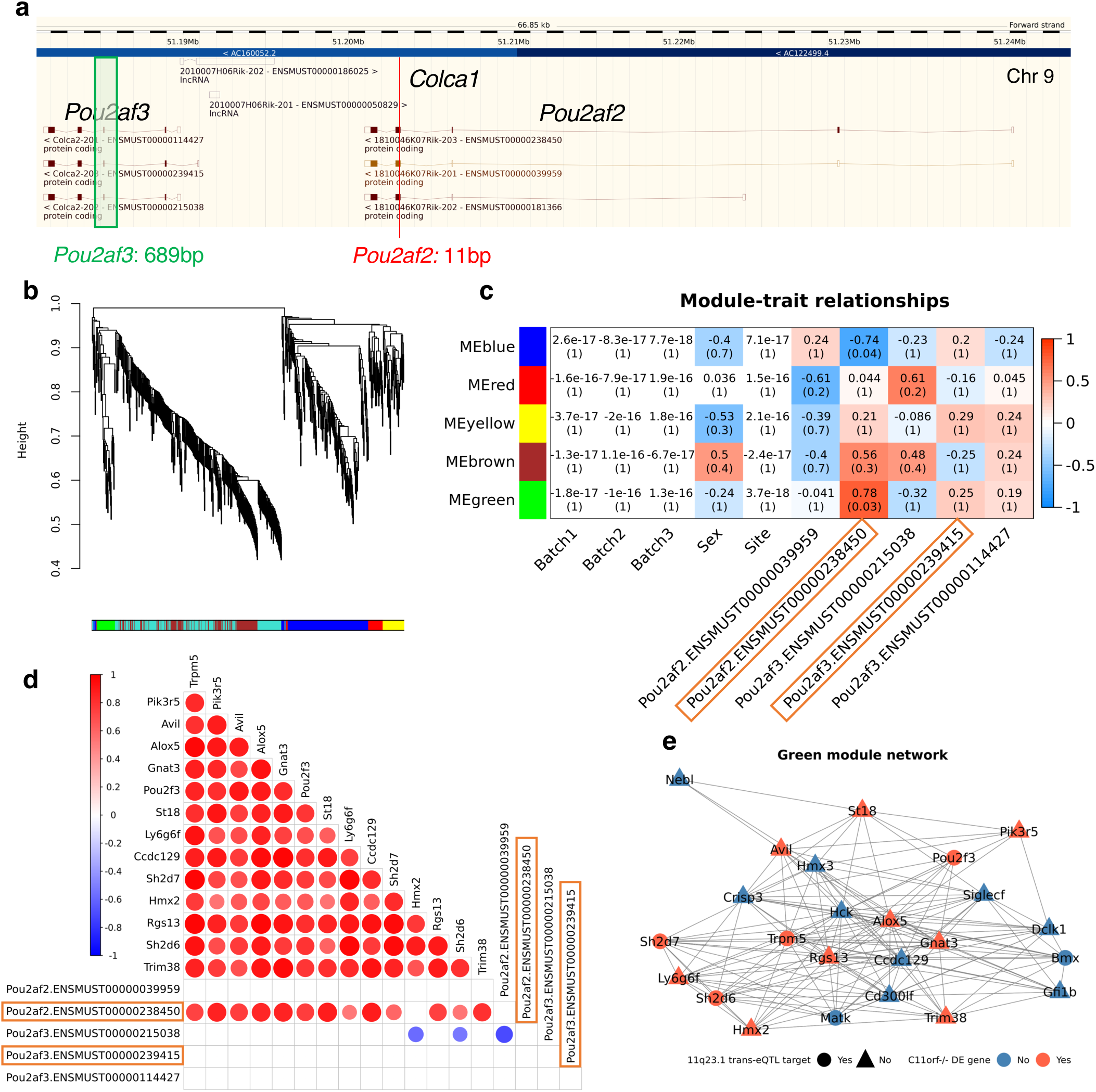
11q23.1 trans-eQTL target homolog expression is correlated with Pou2f interaction domain encoding *Pou2af2* transcript specifically. (a) Generation of the *Pou2af2* and *Pou2af3* models. Schematic obtained from ensembl (https://www.ensembl.org/, accessed 22/09/22). (b) WGCNA of the expression of genes nominally significantly downregulated *C11orfΔ* mice (p<0.01, n=584), across *Pou2af2^-/-^* (n=2) and wild-type (n=7) colon RNAseq. (c) Sample trait matrix of the pearson correlation (above) and FDR-corrected significance (below) between module eigengenes and sample traits, including cis-eQTL target homolog transcripts (boxed if contain POU2F-ID) (d) Pairwise Pearson correlations (p<0.05) between genes downregulated in *C11orfΔ^-/-^* mice (Fig 5b) identified in the green module, in addition to *Pou2af2* and *Pou2af3* transcripts. (e) Kamadakawai network of green module gene relatedness (adjacency >0.3).

With scope to identify *Pou2af2*-regulated genes, we conducted RNAseq analysis of *Pou2af2^-/-^* (n=2) and wild-type (n=7) colonic mucosa. To agnostically test the contribution of individual *Pou2af2* transcripts to 11q23.1 transcriptional dynamics in these samples, we subjected the expression of differentially expressed genes between *C11orf11^-/-^* and wild type mice (p<0.01, n=584) to Weighted Gene Co-expression Network Analysis (WGCNA)^39^, Fig 7b. WGCNA raises gene expression to a power that yields a scale free topology network, then hierarchically clusters genes based on pairwise correlations to identify gene modules that are likely to comprise a co-regulated or co-functional network^39^. The eigengene (first principal component) of the green gene module is highly and significantly correlated with expression POU2F-ID transcript of *Pou2af2* (correlation=0.78, FDR=0.03), but not the non-POU2F-ID *Pou2af2* transcript (correlation=-0.041, FDR=1), or any *Pou2af3* transcripts (maximum correlation=0.25, minimum FDR=1), Fig 7c. The green module includes several genes differentially expressed between *C11orf11^-/-^* and wild-type mice, including; *Trpm5, Pik3r5, Avil, Alox5, Gnat3, Pou2f3, St18, Ly6g6f, Itprid1* (murine homolog of *CCDC129*)*, Sh2d7, Hmx2, Rgs13, Sh2d6* and *Trim38*, indicating reduced expression of these genes in *C11orf11^-/-^* mice is attributable to the expression of the POU2F-ID *Pou2af2* transcript, Supplementary Material 3. In addition, all the previously identified differentially expressed genes present in the green module, excluding *Hmx2,* were correlated with expression of the POU2F-ID *Pou2af2* transcript (p<0.05, median correlation=0.76), but not the non-POU2F-ID *Pou2af2* transcript or any *Pou2af3* transcript (Fig 7d). The hub genes of the green module, as defined by expression adjacency >0.3, also include several 11q23.1 trans-eQTL target homologs; *Trpm5, Pou2f3, Sh2d7*, *Alox5 and Avil*, highlighting them as drivers of this correlation, Fig 7e. Together, this experimentally decouples the expression of 11q23.1 trans-eQTL target homolog to be attributable to *Pou2af2* but not *Pou2af3*, highlighting that this is likely dependent on the POU2F-ID transcript specifically.

In addition, we also assessed the phenotypes found in *C11orf11^-/-^* mice in our single gene knockout models. *Pou2af2^-/-^* mice weighed significantly less than both *Pou2af2^+/-^*(p=8e-3) and wild-type mice (p=4.98e-4) throughout life, which was not observed in *Pou2af3^-/-^* mice, Supplementary Fig S9. In contrast, *Pou2af3^-/-^* males were infertile, a phenotype not observed in *Pou2af2^-/-^* mice, supporting further decoupling of the function of these genes across tissues.

To assess whether the cell-specific expression of 11q23.1 eQTL targets in mice was also analogous to humans, we utilised publicly available scRNAseq of wild-type mouse colonic epithelium^40^. Following dimensionality reduction, we identified five clusters, including one which transcriptionally resembled tuft cells, Fig 8a. Markers of murine tuft cells were significantly enriched for the genes differentially expressed between *C11orf11^-/-^* and wild-type mice (NES=2, FDR=2.3e-03) and green module hub genes previously identified to be correlated with the POU2F-ID *Pou2af2* transcript (NES=2.2, FDR=7.00e-05), indicating transcriptional dynamics of *C11orf11^-/-^* and *Pou2af2^-/-^*mice are attributable to perturbation of this cell-type, Fig 8b. As expected, *Pou2af2* expression is highly specific to a subset of tuft cells, concordant with its identification as a marker of this cell type, Fig 8c and Supplementary Material 5. However, *Pou2af3* expression, while expressed in a minority of tuft cells, shows diffuse expression across goblet cells. As there is evidence to support the shared differentiation trajectory of these cell-types^41^, we analysed the abundance of goblet cells across *Pou2af2* and *Pou2af3* genotype by immunohistochemistry of goblet marker, *Clca1,* and Periodic Acid Schiff (PAS) staining of neutral mucopolysaccharides, present within and secreted by goblet cells, Fig 8d. The abundance of Clca1 was moderately reduced in *Pou2af3^-/-^*mice (p=0.067), but not *Pou2af2^-/-^* mice. Concordantly, we also observed a modest, but not statistically significant, reduction of PAS staining in *Pou2af3^-/-^* mice, but not in *Pou2af2^-/-^* mice. Together, this may indicate the divergent expression of *Pou2af3* to be weakly associated with the differentiation, and potentially function, of murine goblet cells.

**Figure 8.**
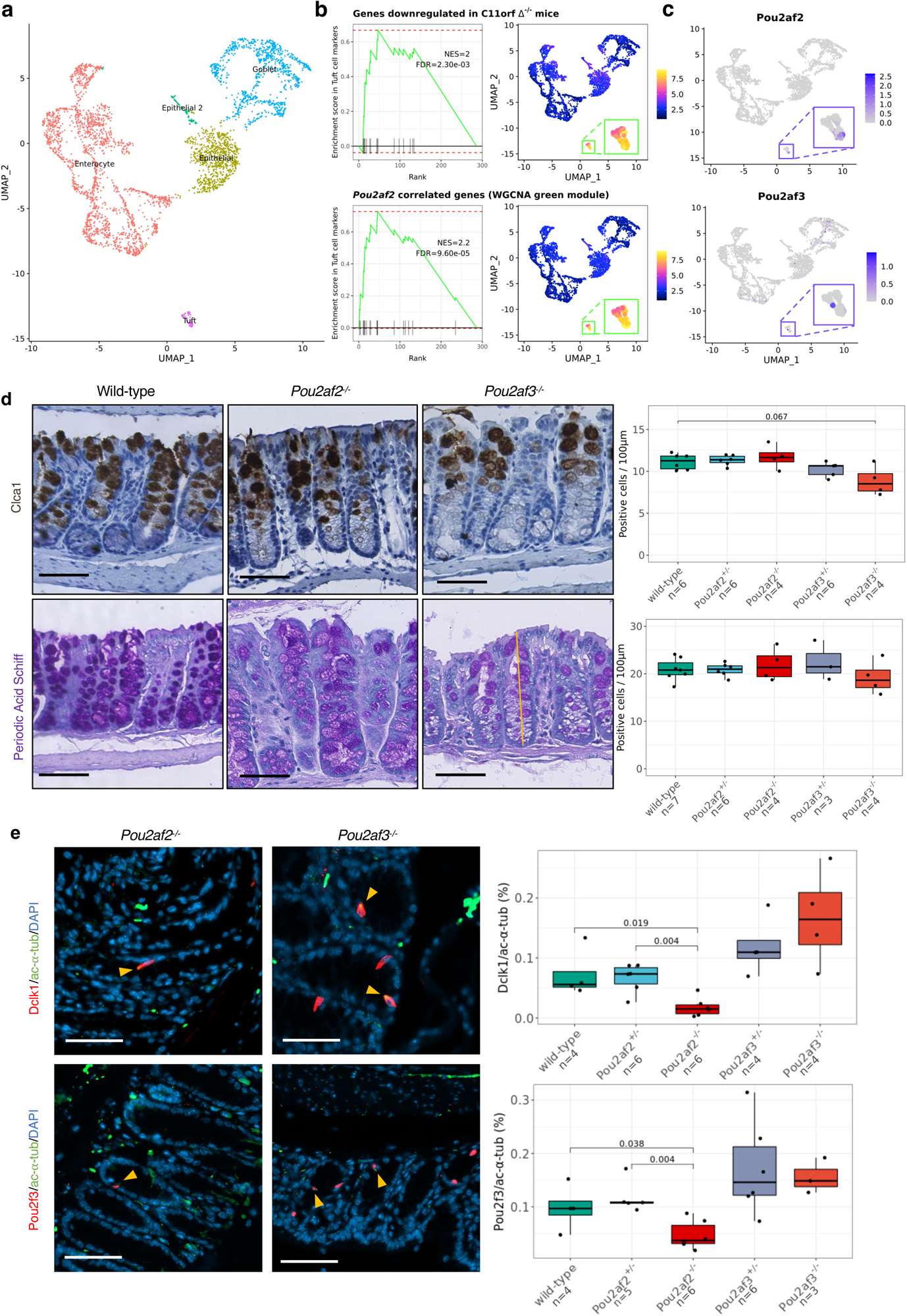
*Pou2af2* and *Pou2af3* expression is divergent across murine colonic epithelium and correlates with the abundance of tuft and goblet cells respectively. (a) UMAP embedding of scRNAseq from 16,828 healthy mouse colon epithelial cells^37^. Cells are annotated by confidently annotated cell-cluster, see Methods. (b/c) GSEA (left) and ssGSEA (right) of *C11orfΔ^-/-^ vs* wild-type differentially expressed genes (top, Fig. 5b) and *Pou2af2* correlated genes from the green module (bottom, Fig. 6c) in murine tuft cell markers and across single cells. (d) Raw expression of *Pou2af2* and *Pou2af3* across individual cells in (a). (e) Abundance of Clca1 and Periodic Acid Schiff stains of goblet cells and mucins respectively, across *Pou2af2* and *Pou2af3* genotypes. P-values are calculated by unpaired Wilcoxon rank sum test. Count number is normalised to the length of crypts along their axis, as shown by yellow bar. (f) Abundance of double positive cells for Dclk1/ac-⍺-tub and Pou2f3/ac-⍺-tub markers of tuft cells by immunofluoresence (Fig 5d) across *Pou2af2* and *Pou2af3* genotype. P-values are Benjamini-Hochberg corrections of unpaired Wilcoxon rank sum test. All scale bars are 50μm.

To assess tuft cell abundance in our single knockout models, we performed immunofluorescence as before (Fig 6e). In accordance with our multi-omic mapping of tuft cell marker expression and accessibility to *POU2AF2* in humans, the abundance of Dclk1/ac-α-tub and Pou2f3/ac-α-tub stained cells was significantly reduced in *Pou2af2^-/-^* (p=0.019, p=0.038 respectively), but not *Pou2af3^-/-^* mice, Fig 8e. Together, this evidence indicates the 11q23.1 variation associated expression changes and reduced colonic tuft cell abundance is a direct result of *Pou2af2* and not *Pou2af3* expression.

#### *Pou2af2* expression is suppressive of colonic tumourigenesis

The association between the expression of CRC risk-associated trans-eQTL target homologs on *Pou2af2* expression, implies; i) 11q23.1 variation may confer CRC risk via dysregulation of this gene specifically, and ii) causal relevance of the associated tuft cell abundance changes. To experimentally interrogate the tumour suppressive effects of *Pou2af2* and *Pou2af3* expression independently, we crossed our single knockout colonies onto a heterozygous *Apc* mutation background that harbours multiple intestinal neoplasia^42,43^, known as *Apc^Min/+^*. *APC* encodes a negative regulator of Wnt signalling; mutated in up to 90% of somatic CRC cases^44,45^, and the perturbed gene predisposes to familial adenomatous polyposis coli. Interestingly, the proportion of *Apc^Min/+^Pou2af2^-/-^*mice generated was less than that expected for mendelian inheritance patterns (chi-squared p=0.06), Supplementary Table S9, potentially implicating *Pou2af2* in exacerbation of Wnt signalling regulation. *Pou2af2* genotype was also significantly associated with morbidity of *Apc^Min/+^* mice (Likelihood Ratio Test=24.14, p=6e-6), where *Apc^Min/+^Pou2af2^-/-^*mice exhibit dramatically reduced survival, Fig 9a. In contrast, *Apc^Min/+^Pou2af3^-/-^*mice were generated at expected frequencies (p=0.19, Supplementary Table S9), and *Pou2af3* genotype was not associated with changes in overall survival on an *Apc^Min/+^* background (LRT=0.62, p=0.7).

**Figure 9.**
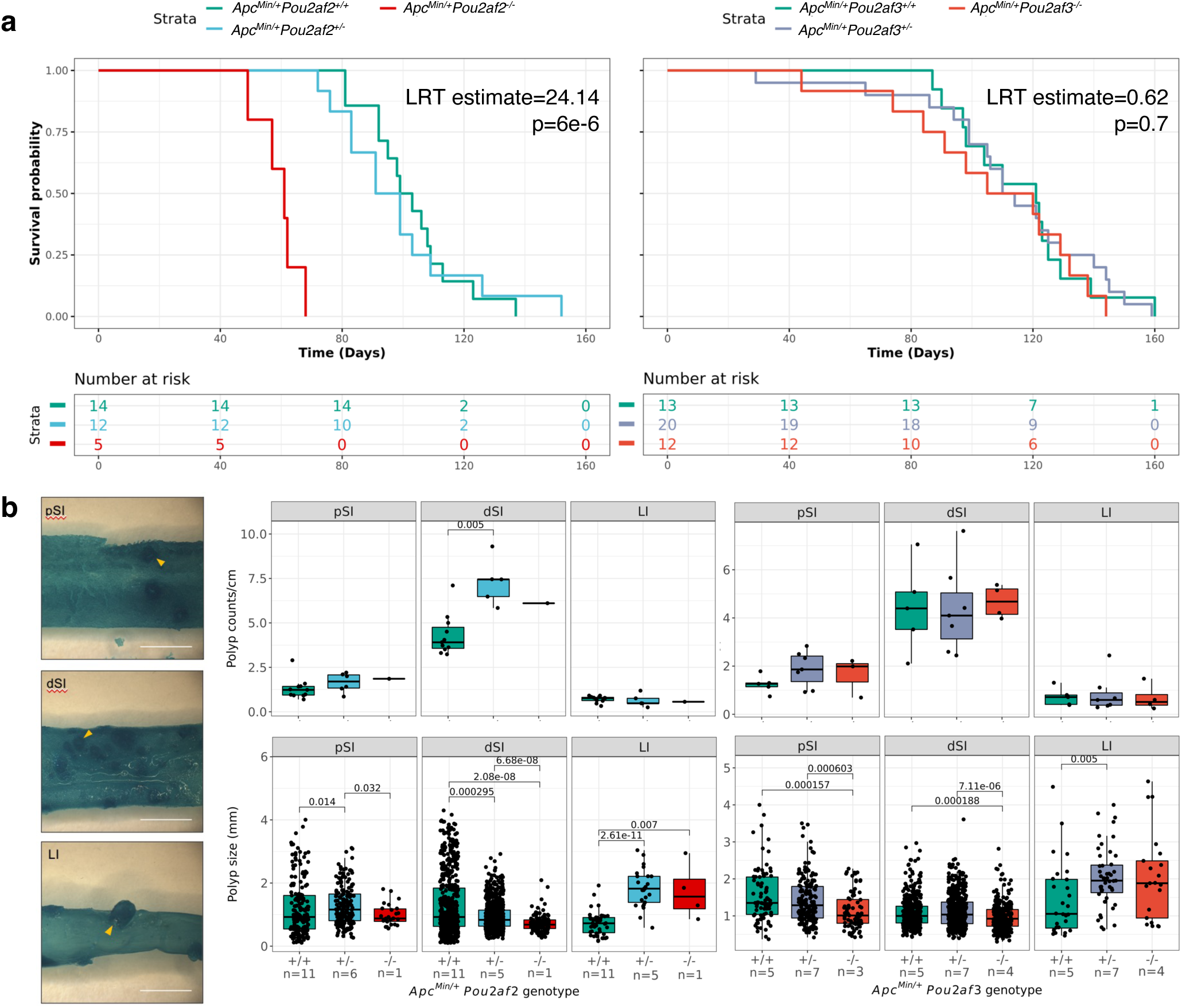
Expression of *Pou2af2,* but not *Pou2af3,* is protective of tumourigenesis in *Apc^min/+^* mice. (a) Kaplan-Meir survival analysis of *Apc^Min/+^Pou2af2* (left) and *Apc^Min/+^Pou2af3* (right) mice, LRT=Likelihood ratio test. (b) (Left) Example images of methylene blue stained intestinal samples, with yellow arrowhead identifying example polyps. Scale bars are 1cm. (Right) Abundance and size of methylene blue-stained polyps across *Apc^Min/+^Pou2af2* and *Apc^Min/+^Pou2af3* mice (middle and right). pSI=proximal small intestine, dSI=distal small intestine, LI=large intestine/colon. P-values are Benjamini-Hochberg corrections of unpaired Wilcoxon rank sum test.

To investigate whether the altered survival of *Apc^Min/+^* mice across *Pou2af2* genotype was specifically associated with intestinal tumourigenesis, we analysed polyp frequency and size by staining small intestine and colon samples with methylene blue, a well characterised stain for aberrant crypts and therefore intestinal polyps^46^, Fig 9b. Unfortunately, this analysis is limited to small numbers of *Apc^Min/+^Pou2af2^-/-^* mice due to inability to consistently generate these mice (Supplementary Table S9), conformation with NC3R ARRIVE guidelines^47^ and inability to collect samples due to COVID restrictions. However, *Apc^Min/+^Pou2af2^+/-^* mice exhibited an ∼2-fold increase in polyp number in the distal small intestine compared with *Apc^Min/+^* mice (FDR=0.005), in addition to increased polyp size in the proximal small intestine (FDR=0.014) and colon (FDR=2.61e-11). Similarly, the *Apc^Min/+^Pou2af2^-/-^* mouse exhibits significantly larger polyps in the colon than *Apc^Min/+^* mice (FDR=0.01), further supporting *Pou2af2* to be associated with increased tumour initiation and progression on this background.

In contrast, *Pou2af3* genotype was not associated with any significant changes in tumour frequency in the small intestine or colon of *Apc^Min/+^*mice, Fig 9b. Polyps were also significantly smaller in *Apc^Min/+^Pou2af3^+/-^*and *Apc^Min/+^Pou2af3^-/-^* proximal small intestines (FDR=1.57e-4, FDR=8.04e-4 respectively) and distal small intestines (FDR=2.5e-4, FDR=7.11e-6 respectively). While there was a small increase in the size of colonic polyps from *Apc^Min/+^ Pou2af3^+/-^* mice (FDR=0.005), this was not significantly increased in *Apc^Min/+^Pou2af3^-/-^* mice. Taken together, and in the context of exclusively *Pou2af2-*associated survival changes, this shows downregulation of *Pou2af2* is associated with exacerbated tumour initiation and progression in this model.

## Discussion

In this study, we robustly identify several 11q23.1 trans-eQTL targets, confirm their relevance to CRC risk, and delineate their genetic association to a single signal at rs3087967. Present and previous^14^ genome-wide, molecular characterisation of healthy human colonic epithelium at a single-cell resolution strongly implicates *POU2AF2* as a regulator of the tuft cell-specific expression of 11q23.1 trans-eQTL targets, a finding we experimentally delineate to *Pou2af2* across multiple knockout mouse models. We also confirm CRC risk genotype at 11q23.1 to correlate with reduced tuft cell abundance in the human colon, and assess the contribution of *Pou2af2* and *Pou2af3* expression to tumourigenesis in mice, defining a tumour suppressive effect of *Pou2af2* expression and by correlation, colonic tuft cell abundance.

Convergence of CRC risk and cis-eQTL effects to a single signal at rs3087967 both reinforces casual relevance of cis-eQTL effects and implies this site in the underlying mechanism of gene dysregulation. In addition, as the trans-eQTL targets identified in the transverse colon were not found in any other GTEx site and 11q23.1 variation associated cancer risk has not been identified in any other tissue, this also suggests relevance of these transcriptional dynamics in governing CRC risk. Importantly, we statistically confirm causal relevance of 11q23.1 trans-eQTL targets by integration with the largest CRC risk GWAS to date^6^, identifying a myriad of novel CRC risk-associated genes. Interestingly, the lack of cis-eQTL effects at these regions supports their expression to be predominantly regulated by 11q23.1 genetic variation effects, which is in complete concordance with the transcriptionally activating role of POU2AF2 identified for these genes by ChIPseq analysis. While the exact mechanism of cis-eQTL target dysregulation remains to be elucidated, as rs3807967 lies within the 3’ UTR of *POU2AF2,* this suggests it may occur through altered transcript stability, also reinforcing altered expression of *POU2AF2* as the primary mechanism by which cis-eQTL target dysregulation occurs. Of course, assessment of transcriptional changes through a genetic screen of 11q23.1 eQTLs with functional expression readout would provide a gold-standard identification of the causal variant at this region, and likely provide strong indication of the dysregulatory mechanism.

While our recent work potentiated de-coupling of cis-eQTL association with trans-eQTL target expression^14^, until the present study, this remained to be experimentally validated. We identified non-congruent chromatin accessibility profiles of *POU2AF2* with *COLCA1* and *POU2AF3,* conforming with divergence of their genetic regulation in the human colon. We also identified likely transcriptionally activating genomic binding of POU2AF2 at 11q23.1 trans-eQTL targets, the majority of which were selectively accessible in colonic tuft cells. While POU2AF3 ChIPseq was not available, our genetically engineered mouse models, which accurately recapitulate 11q23.1 transcriptionally dynamics, experimentally delineate the transcriptional dynamics and tuft cell abundance to *Pou2af2* over *Pou2af3*, concordant with a transcriptionally activating role of *POU2AF2* in colonic tuft cell development. Our delineation of this effect to *POU2AF2* does contrast findings by Wu *et al.,*^19^, which imply *POU2AF3* expression to be associated with tuft cell marker expression in SCLC-P models. It is possible that fine-tuning of tuft cell abundance by *POU2AF2* and *POU2AF3* varies across tissues, and because a weak cis-eQTL signal is detected for *POU2AF2* but not *POU2AF3* in the lung, with no preservation of trans-eQTL effects, it is likely regulation of tuft cell abundance in this tissue differs to that of the colon. However, because the expression of *POU2AF3* is dramatically less well correlated with 11q23.1 trans-eQTL targets in human colonic tuft cells, is not highly expressed in murine colonic tuft cells, and was not associated with the abundance of murine colonic tuft cells in our models, we are confident colonic tuft cell abundance is primarily associated with *POU2AF2* expression.

Tuft cells are associated with paracrine stem-, neurotransmitting- and immune-related functions^15,48–51^, including a well-characterised mechanism of defence against helminth infection^52–54^. However, much of this evidence is derived from the small intestine and cannot necessarily be extrapolated to the colon. The role of tuft cell markers in governing such processes has been well documented^55,56^ and while we do not investigate the functions of these proteins directly, we experimentally determine reduced tuft cell abundance to correlate with CRC risk in humans. Accordingly, reduced tuft cell abundance has been associated with pancreatic tumourigenesis in independent studies^57,58^. In both cases, this occurs through alteration of the immune related signalling, implicating altered immune regulation as a potential route by which CRC risk occurs in the colon. In addition, lower abundance of tuft cells has been observed in the colon of quiescent ulcerative colitis patients^59^. As there is a well-known association between colitis and CRC risk^60–62^, reduced tuft cell abundance may be relevant in colitis-associated CRC (CA-CRC) also. The potential relevance of 11q23.1 genes in CA-CRC is compounded by genetic mapping studies identifying the homologous region in mice to mediate susceptibility of mouse models to CA-CRC^63^. Notably, recent work has also implicated TET2/3 as critical mediators of the response of mice to chemically induced colitis by regulation of *POU2F3* methylation and small intestinal tuft cell abundance^64^, further supporting the importance of this cell-type in governing immune response in the gut. While our study does not provide evidence of the method by which tuft cell-associated functions may yield potential CRC risk in the colon, it should remain to be a focus of future work.

To our knowledge, this is the first study to identify *POU2AF2* or any other cis-eQTL target for any disease loci, to be associated with heritable changes of cell abundance in the colon. In addition, our concordant experimental delineation of 11q23.1 variation associated tumourigenic risk to *Pou2af2* over *Pou2af3* is of great interest. While it is possible that independent *POU2AF3* or *COLCA1-*CRC risk associated mechanisms exist, our study provides strong evidence that the reduced tuft cell abundance conferred by *POU2AF2* is correlated with CRC risk. Further characterisation of this and other CRC genetic risk-associated mechanisms is likely to uncover potential preventative, therapeutic or patient stratification mechanisms that reduce disease prevalence, mortality and burden.

## Materials and Methods

### Ethics statement

scRNAseq^65^ and scATACseq^32^ of healthy colonic mucosa are previously published. For human tissue used in this study (RNAseq and immunofluorescence), all participants provided informed written consent, and research was approved by local research ethics committees (SOCCS 11/SS/0109 and 01/0/05; SCOVIDS 13/SS/0248) and National Health Service management (SOCCS 2013/0014, 2003/W/GEN/05; SCOVIDS 2014/0058).

### Data and code availability

Publicly available datasets utilised in this study and their corresponding access codes are summarised in Table S10. The GTEx v8 data used for the analyses described in this manuscript were obtained from the GTEx Portal on 11/05/21 and dbGaP accession number 23765. For Generation Scotland (GS) data access is through the GS access committee (GSAC). Applications for the Lothian Birth Cohort data should be made through https://www.ed.ac.uk/lothian-birth-cohorts/data-access-collaboration. Code used to perform all analysis of this study is available at https://github.com/BradleyH017/.

### RNA extraction and qRT-PCR of human peripheral blood and colorectal mucosa

For human normal mucosa and peripheral blood, RNA was purified using RiboPure RNA purification kit (Ambion) according to manufacturer’s protocol. Purified RNA was treated with DNase (Promega) at 37°C for 30 min. To convert DNase treated RNA into cDNA, 1μl of random primers (Promega), RNasin Plus RNase inhibitor (Promega), dH_2_O and M-MLV Reverse Transcriptase (Promega) was added to RNA, along with 2μl of dNTPs (10mM Invitrogen) and 4μl of 5X M-MLV reaction Buffer (Promega). Samples were then incubated at 37°C for 1 hour, followed by 95°C for 10 mins. To assess gene expression, we utilised Taqman probes summarised in Table S11. Reactions contained 0.5μl of probe (Applied Biosystems), 2.5μl DNase/RNase free water and 2μl cDNA per well, of which were completed in triplicate. The reaction was amplified and detected on an ABI PRISM 7900HT detector system with conditions of 50°C for 2 mins, 95°C for 10 mins followed by 40 cycles of 95°C for 15 seconds and 60°C for 1 minute. Data was analysed using SDS software (Applied Biosystems). Gene expression in samples was calculated using linear regression analysis of the standard curves. Gene expression was also normalised to an endogenously expressed gene. Gene expression values was normalised to *TBP*, *EIF2B1* and *RPL30* (normal mucosa) or *ACTB* (blood). All qRT-PCR data is presented as log base 2 of normalised expression values.

### Fine mapping and eQTL analysis

For each GTEx site analysed, the expression data was subset for those with corresponding whole genome sequence data available. Variants were subset for those which had a minor allele frequency of >0.01 within the subset data and located within 1Mb 5’ of *POU2AF2* and 1Mb 3’ of *POU2AF3*. As per GTEx quality control, the expression data was also subset to remove lowly expressed genes by intersection of genes that exhibited i) >0.1 transcripts per million in at least 20% of samples, ii) >= 6 reads in 20% of samples. Expression was subsequently inverse normally transformed using *RNOmni* v1.0.0 ^66^ before hidden covariates were identified using a maximum of 10,000 model iterations by *PEER* v1.0 ^67^. Binarized sequencing batch, binarized sex and age were accounted for, in addition to a number of hidden covariates equivalent to a quarter the number of samples for each site. Residual expression was subsequently re-normally distributed and used to test association between variants and genome-wide expression using *MatrixEQTL* v2.3^68^. P-value output thresholds for both local and distant associations were 0.01. The same procedure was followed for analysis at the transcript level, and FDR values incorporate genome-wide multiple testing comparisons. Manhattan plots were generated using *locusZoom* v0.12^69^ web interface.

### Colocalization analysis

Colocalization analysis on gene expression and CRC risk was performed with the *coloc* package v5.2.1^70^. CRC risk GWAS summary statistics for a recent GWAS^6^ were obtained from GWAS catalog using accession GCST90129505. Analysis was done to assess the relative significance of global hypotheses, using a p12 parameter of 1e-4. To identify variants likely associated with the common, single variant hypothesis, a 95% credible set of variants was subset from the results.

### Conditional analysis

To perform conditional analyses, we utilised *gcta* v1.91.4beta^71^ software in –cojo-actual-geno mode, conditioning on rs3087967 and accounting for distribution and frequency of variants tested in the GTEx population (for conditioning of expression) or 1000G European reference panel (for conditioning of CRC risk).

### Summarised Mendelian Randomisation

Trans-eQTL effects (>1Mb) of 11q23.1 were calculated for whole genome sequencing variants within 1Mb of 11q23.1 cis-eQTL targets in GTEx transverse colon (n=367) and SOCCS (n=213) as before, see *Fine Mapping and eQTL analysis.* The summary statistics from this were then merged using a fixed effects model, performed using *META* v1.7^72^. Cis-eQTLs were also detected within 1Mb of trans-eQTL target genes, and meta-analysed across datasets in the same manner. SMR was then performed to assess the association between gene expression and CRC risk using *smr* v1.3.1^30^, recently published CRC risk GWAS summary statistics^6^ and the 1000 genomes plus UK10,000 genomes reference panel. To protect against artificial inflation of SMR significance by focussing on 11q23.1, 11q23.1 trans-eQTLs were only utilised for SMR if their significance was greater than any cis-eQTL detected for that gene.

### CRC risk interaction analysis

Genotype data was obtained for three studies, namely Generation Scotland, the Lothian Birth Cohort and the National Study of Colorectal Cancer Genetics, totalling 30,556 individuals. Details on imputation and QC have been previously published^5^. The interaction analysis was performed using plink v1.90b6.18 using the epistasis function. The alleles of the lead variant at the 11q23 locus (rs3087967) were compared to all variants within 1Mb of significant eQTLs. Significance was determined using logistic regression in the case-control analysis, and a Chi-square test in the case only analysis. Meta-analysis of the results was performed using the R meta package.

### Transcript encoding isoform analysis

GRCh38 transcript cDNA sequences of *POU2AF2* and *POU2AF2* were first obtained from ensembl (https://www.ensembl.org/index.html). To convert into protein sequence, the cDNA from the first AUG start was input into EMBOSS Transseq (https://www.ebi.ac.uk/Tools/st/emboss_transeq/). POU2F interaction domain encoding was then identified, limiting search from the first amino acid to the first stop codon.

### Human scRNAseq analysis

Dimensionality reduction and clustering analysis of human colon scRNAseq data^65^ was performed as previously described^14^. To compute correlation of gene expression at the single cell level, we extracted the expression matrix and imputed technical dropout events using *Rmagic* v2.0.3^73^. Pearson correlations were only computed for genes within clusters when at least 50% of comprising cells had imputed expression scores >0.01. Pearson correlations were only included in analysis if they passed correlation significance p<1E-3 and coefficient >0.4.

### Mouse scRNAseq analysis

Wild-type mouse colonic epithelial scRNAseq generated by Tabula Muris Consortium^40^ was obtained from gene expression omnibus using accession GSE109774. Counts were pre-processed to remove bad quality cells as previously described^14^. The filtered count matrix (16,828 genes and 3,853 cells) was loaded into a Seurat object using *Seurat* v4.1.1^74^. Data was normalised using SCTransform before finding variable features, PCA calculation and UMAP dimensionality reduction using the first 20 principal components. The kNN graph was calculated using 20 nearest neighbours and cells were clustered using a resolution of 0.6. A log transcript per 10,000 gene count matrix was also calculated as previously described^14^.

For cell annotation, a second dataset was obtained from PanglaoDB using accession number (SRA653146). The authors cell annotations were used to calculate cluster markers based on log-normalised expression values. Gene set enrichment was then performed on our analysis of Tabula Muris data against the second dataset using *fgsea* v1.16.0^75^, annotating clusters significantly enriched for annotation in the second dataset (FDR<0.05) or manual inspection, resulting in the 4 confidently annotated clusters (Tuft, Goblet, Enterocyte, Epithelial) and one remaining cluster, ‘Epithelial 2’. Enrichment of signatures at the single cell level was performed using *escape* v1.0.0^76^.

### ChIPseq analysis and motif enrichment

Narrowpeak files of POU2AF2 and POU2F3 ChIPseq results, along with POU2AF2, POU2F3, p300, MED1 and acetylated H3K27 bigwig files were downloaded from gene expression omnibus using accession code GSE186614^19^. These were annotated on the hg38 genome using *annotatePeaks.pl* and default parameters from *HOMER* v4.1.1^77^. POU2AF2 binding was then identified in each narrowpeak file and saved as a bed file, combining both cell lines and antibody replicates. Motif discovery was subsequently executed using *findMotifsGenome.pl* on the bed file, with fragment size parameter set at 200 bases. To identify motif presence at each refined 11q23.1 trans-eQTL target, sequences of POU2AF2-bound regions were obtained using *Biostrings* v2.60.0^78^, and bound sequences were searched for ATTTGCA/TTTGCAT (POU2F3), ATGCAAAT (Oct2) or ATTTGCAT (Oct4). Read enrichments of POU2F3 and POU2AF2 binding were plotted from the authors bigwig files using the *wiggleplotr* v1.18.0 package^79^.

### scATACseq analysis

Bed files from scATACseq of healthy colonic normal mucosa^32^ were obtained from Gene Expression Omnibus using accession GSE184462. This was first subset for the colon epithelial samples by intersecting with cell-level metadata from this study, available on catlas (http://catlas.org/humanenhancer/#!/cellBrowser). All subsequent analysis was performed using the archR analysis pipeline and management package, v1.0.1^80^; i) an iterative LSI dimensionality reduction was generated using the tile matrix, ii) batch correction was performed on the Iterative LSI matrix across samples using Harmony v1.0^81^, iii) Clusters were identified using the ‘Seurat’ method and 10 nearest neighbours on the Harmony corrected matrix, iv) UMAP embeddings were calculated on the Harmony corrected matrix, v) Accessibility around entire gene regions was calculated and used to determine cluster markers, accounting for transcription start state enrichment and log10(Fragment number), vi) Enrichment of POU2AF2 bound regions was performed by calculating peak annotation of the narrowpeak files for each antibody and cell-line replicate (see *ChIPseq analysis and motif enrichment*), vi) Mean accessibility of POU2AF2 bound regions within each cluster were manually calculated after extracting the POU2AF2 accessibility deviation matrix, vi) P-values for changes in accessibility across clusters were calculated by t-test of the accessibility in cluster 5 compared to all other clusters combined.

### Human colonic epithelium collection and processing

Following bowel resection, epithelium was removed from the full thickness mucosa using Metzenbaum scissors, shown in Supplementary Video S1. Samples were subsequently fixed in 10% neutral buffered formalin for 24 hours at room temperature and stored in 70% ethanol at 4°C. To perform swiss-rolling, colonic tissue strips were first lay on foil-covered polystyrene with the epithelium side facing down. One end was pinned using two 25G needles and the entire sample was lay flat and taught by pinning with additional 25G needles. Samples were then rolled around a toothpick towards the initially pinned needles, removing the additional needles as they were met. Samples were then paraffin embedded using a single 25G needle to maintain roll structure and cut cross sections at 5μm thickness. An example of the swiss-rolling method is shown in Supplementary Video S2.

### Genotyping of human samples

Patient blood samples were acquired ahead of surgery, Supplementary Table S12. DNA was extracted using Qiagen blood and tissue extraction kit (#69504), as per manufacturers protocol for non-nucleated blood. To genotype samples at rs3087967, 1.5μl of extracted DNA was added to 1.25μl 20μM forward primer (TGGAAGATCTGCACCACACT), 1.25μl 20μM reverse primer (ATGCCCTCGTCCACTAACAA), 25μl of DreamTaq Green PCR MasterMix (ThermoScientific #K1081) and 21μl of dH_2_O. Amplification was performed by heating to 95°C for 5 mins, followed by 35 rounds of 95°C for 30 seconds, 55°C for 30 seconds and 72°C for 30 seconds and a final incubation at 72°C for 5 mins. The PCR product was subsequently sanger sequenced by the IGC Technical Services facility and rs3087967 genotype was analysed using ApE software v2.0^82^.

### Genetic perturbation of mice

All animal studies were approved by the University of Edinburgh ethics committee and performed under a UK Home Office project license.

#### CRISPR guide generation

Plasmid px458 was digested with restriction endonuclease BbsI (New England Biolabs, R0539S), by addition of 1μl enzyme to 1μg plasmid, 0.2μl 100X Bovine Serum Albumin (New England Biolabs, B9000S), 200μl 10X NEBuffer 2.1 (New England Biolabs, B7202S) and dH_2_O to a final volume of 20μl. The digestion reaction was then incubated at 37°C for 70 mins. Guide RNA (gRNA) sequences (Supplementary Table S14) were purchased as single stranded DNA oligonucleotides, together with reverse complements. Oligonucleotides were annealed by combining 5μl of each primer at 100μM, along with 10μl of dH_2_O and incubated for 37°C for 30 mins, 95°C for 5 mins and cooled to 10°C at 0.1°C/second. 1μl of diluted (1:200), annealed oligonucleotides was ligated into linearised px458 by incubation with 0.5μl digested plasmid, 2.5μl dH_2_O, 5μl 2X T4 ligase buffer (New England Biolabs, B0202S) and 1μl T4 DNA ligase (New England Biolabs, M0202S) at room temperature for 30 mins. Plasmids were transformed into competent cells by mixing with 2.5 μl ligation mixture and incubation for 30s on ice, 42°C for 30s and 5 mins on ice. 950μl SOC medium (Thermo Fisher Scientific, 15544034) was added to each tube and incubated on shaker in 37°C incubator for 1 hour. Each culture was plated on two warm L-amp plates (IGC Technical Services). Cultures were spread around whole plate using cell spreader. Plates were incubated at 37°C overnight and examined for colonies. Colonies were picked into 5ml L-broth with ampicillin. Cultures were incubated on shaker in 37°C incubator overnight. Plasmids were extracted from expanded colonies using the QIAprep Spin Miniprep kit (Qiagen, 27104), according to supplied kit instructions.

#### Injection into mice

Extracted plasmids were defrosted on ice. For each gRNA combination, an injection mix was prepared including 50ng/μl Cas9 mRNA (Tebu Bio, L-6125-20), 25ng/μl each gRNA, 150ng/μl each repair template, dH2O to total volume 50μl. These were injected into C57/Bl6 embryos by BRF/Evans transgenic unit.

The *Pou2af3* mouse model (alternative name: Gm684_tm1c_C08, code: ABF, Strain Name: C57BL/6N-Colca2/Tcp, MGI:7257808) was obtained from the Canadian Mutant Mouse Repository (Bradley *et al.,* 2012).

### Dissection of mouse intestinal tissue

Mice were culled by cervical dislocation. For dissection, animals were pinned on their back and an incision made along ventral midline. Small intestine samples were collected from duodenum up to but not including caecum and divided in half to produce the proximal and distal fractions. The colon was taken from (but not including) the caecum to (and including) the rectum. Samples were washed with PBS and inverted on skewers before fixation in 10% neutral buffered formalin for 24 hours at room temperature. Following fixation, these were rolled along the length of the sample using a toothpick and paraffin embedded. For staining, slides were cut at 5μm thickness. For RNA collection, 1cm of PBS-washed tissue was collected from the proximal end of each fraction, including a further division of the colon into proximal and distal by dividing in half along the length of the sample from caecum to rectum. Samples were stored in RNAlater (Invitrogen) until extraction.

### RNA extraction and sequencing of mouse colonic mucosa

For mouse tissue, RNA was extracted using phenol-chloroform extraction method and sequenced by polyA selection using a NextSeq machine at 25 million paired-end reads by the Wellcome Trust Clinical Research Facility.

### Bulk RNAseq and WGCNA analysis of mouse colonic mucosa

Raw sequencing reads were aligned to the mm10 genome using Nextflow v21.04.2 nf-core RNAseq sequence alignment pipeline^83,84^. Gene and transcript level counts were subset to remove lowly expressed genes as before *(see Fine Mapping and eQTL analysis*). For differential expression analysis; sex, site and batch were included in the design matrix that was utilised to perform differential expression by quasi-likelihood F test with *edgeR* v3.34.1^85^.

For network analysis, counts were further z-scored before removal of batch and site effects using *limma* v3.48.3^86^ *RemoveBatchEffect*. These were then subset for genes differentially expressed between *C11orf11^-/-^* and wild-type mice and used as input into *WGCNA* v1.69^39^. The recommended scale free topology threshold was identified as 10 and utilised for downstream analysis.

### Immunofluorescent staining of colonic tissue

To deparaffinise the tissue, samples were incubated in xylene for 15 mins before reduced dilutions of ethanol (100%, 90%, 70%) for 10 mins each. Slides were then dipped into de-ionised water (dH_2_O) before 0.1% Tween in PBS for 5 mins. Antigen retrieval was performed by pressure cooker incubation of slides in 1L of pre-warmed 1X Citrate Buffer (Sigma Aldrich #C9999) + 0.1% Tween in dH_2_O for a total of 15 mins (human) or 10 mins (mouse). Slides were then cooled at room temperature for 1 hour after addition of 1L of dH_2_O. Samples were then transferred into a sequenza slide rack (ThermoScientific #73310017) in coverplates (ThermoScientific #72110017) and washed with PBS-T to check for adherence before permeabilization with 500μl of 0.5% Triton-X-100 (BioXtra #T928) for 20 mins at room temperature. Slides were then washed with PBS-T once more before blocking for 1 hour at room temperature with 10% goat serum (abcam #ab7481) or 1% Bovine Serum Albumin (Sigma #05482) in PBS-T for mouse and human samples respectively. 300μl of primary antibody was subsequently added to the slides and left at 4°C for 24 hours. All antibodies and concentrations used in this study are summarised in Supplementary Table S15.

Following primary antibody incubation, slides were washed with PBS-T three times. 300μl of secondary antibody was then added to slides and incubated at room temperature for 2 hours. Slides were then washed with PBS-T 5 times before DAPI stained and mounted with one drop of VectaShield Antifade mounting medium with DAPI (Vectashield, #H-1200).

To analyse immunofluorescent staining, slides were scanned at 20X magnification on Zeiss Axioscan.Z1 using Zen Blue software v3.389 (https://www.zeiss.com/microscopy/en/products/software/zeiss-zen.html). As only double-positive cells were deemed as positive, these were counted blindly by eye in a single swiss-roll only. Background cell number was calculated using the ‘Cell detection’ feature of QuPath v0.2.3^87^ and default parameters, after colour balance removal of red and green signal. For the background cell number, muscle, fat or skin nuclei were not included. For comparison of cell number versus non-swiss rolled method, the background epithelial/stromal cell number for non-swiss rolled tissue was calculated as before, and both swiss-rolled and non-swiss rolled counts were normalised to the squared footprint area the scanned tissue occupied.

### Immunohistochemical staining of mouse colon

Deparaffinisation was performed as before (see *Immunofluorescent staining of colonic tissue*). Before first incubation in PBS-T, slides were kept in 3% hydrogen peroxide in PBS for 20 mins at room temperature. After incubation with the secondary antibody (Supplementary Table 10), nuclei were stained by addition of 300μl DAB solution (Vector #SK-4100), as per manufacturers protocol, and left at room temperature for 5 mins. Slides were subsequently removed from the sequenza and transferred to dH_2_O for 5 mins. Slides were then placed in Harris Haemotoxylin (Sigma-Aldrich #HHS32) for 30s, before being washed with tap water for 5 mins. Samples were then placed in increasing concentrations of ethanol (70%, 90%, 100%) for 10 mins each, before xylene for 15 mins. Finally, slide covers were mounted using Cytoseal XYL mounting medium (ThermoScientific #8312-4).

Images were obtained using a Hamamatsu Nanozoomer XR and analysed using QuPath v0.2.3^87^. Positive cells were counted in each of 40 crypts across one entire colon swiss roll by eye and normalised to the length of each crypt along it’s axis (base to peak).

### Periodic Acid Schiff staining of mouse intestinal tissue

Slides were deparaffinised and hydrated as before (see *Immunofluorescent staining of colonic tissue*). The sections were subsequently placed in periodic acid solution (1%) for 5 mins before being rinsed with distilled water. The slides were then covered with Schiff’s reagent for 20 mins and rinsed under tap water for 5 mins. Counterstaining was performed using Harris haemotoxylin (Sigma-Aldrich #HHS32), before being dehydrated by incubating in 70%, 90% and 100% ethanol for 10 mins each. Finally, slides were incubated in xylene for 15 mins before mounting (see *Immunohistochemical staining of mouse colon*).

### Apc^Min/+^ survival and tumour burden analysis

*Pou2af2* and *Pou2af3* mice were crossed onto a C57Bl6 *Apc^min/+^* strain and aged until moribund phenotypes were observed, defined as; pale paws, rectal bleeding (most common), hunched posture, starry coat or rectal prolapse. Survival analysis was carried out using *survival* v3.3.1^88^, to compute univariate likelihood ratio test estimates and p-values. Hazard ratio tests for *Apc^min/+^Pou2af2^-/-^*exceeded the statistically plausible threshold due to effect size. Univariate tests showed no interaction between sex and survival.

Intestinal samples were stained by dipping into 0.2% methylene blue solution stock 5-6x in dH_2_O (Sigma-Aldrich, #M9140). Tissues were then allowed to de-stain for at least 1 week by incubating in 70% ethanol, changed every 2-3 days before being imaged at 0.8x magnification on a stereomicroscope. Polyp size was measured as the diameter along the same axis as tissue sample and analysis was performed using Image J version 1.52a^89^.

## Acknowledgements

We would like to first acknowledge the excellent management, care and help of Natasha Ackerman and the rest of the technical staff at the University of Edinburgh BRF/Evans mouse facility. In addition, we would also like to thank the sample processing of the Pathology unit at the Institute of Genetics & Cancer, for their tireless and excellent processing of human and mouse tissues and the Genetics Core at the Wellcome Trust Clinical Research Facility. We would also like to acknowledge the collection of blood samples from patients by Donna Markie, which was paramount to genotyping ahead of surgery - critical to targeted sample collection. Finally, we would like to offer great appreciation to all the donors and their families, without whom this work would not have been possible.

## Author contributions

Conceptualisation: V.R., B.T.H., R.T.O., C.S., L.Y.O., M.G.D., S.M.F. Methodology: V.R., B.T.H., R.T.O., C.S., K.D., M.T., S.M.F., M.G.D. qRT-PCR: L.Y.O., C.S. RNAseq read alignment: G.G., A.M., K.D., J.P.B., B.T.H. Fine mapping, conditional, colocalization, summarised mendelian randomisation, scRNAseq, ChIP-seq, scATACseq, differential expression, WGCNA, isoform and survival analysis: B.T.H. CRC risk analysis: P.J.L. Human risk variant genotyping: B.T.H., S.R., M.W. Human colon tissue collection: M.A., M.G.D, F.V.N.D. Human colon tissue processing: V.R., B.T.H. Mouse model generation: V.R., R.T.O. Mouse model maintenance and dissection: V.R., B.T.H., R.T.O., M.B. Mouse phenotyping: V.R., B.T.H., R.T.O., E.E.B., M.B. Immunofluoresence: B.T.H. Immunohistochemistry: B.T.H. Original draft writing: B.T.H. Review & Editing: V.R., B.T.H., M.G.D., S.M.F. Funding Acquisition: S.M.F., I.T., M.G.D., F.V.N.D. Supervision: V.R., R.H., M.G.D., S.M.F. All authors read and approved the final manuscript.

## Open access

For the purpose of open access, the author has applied a CC-BY public copyright licence to any Author Accepted Manuscript version arising from this submission.

## Abbreviations

CRC: Colorectal cancer
GWAS: Genome-wide association studies
RNAseq: RNA sequencing
SNP: single nucleotide polymorphism
scRNAseq: Single-cell RNA sequencing
POU2F-ID: POU2f interaction domain
ChIPseq: Chromatin immunoprecipitation sequencing
scATACseq: Single-cell assay for transposable element accessibility sequencing

---

**Supplementary Figure S1.**
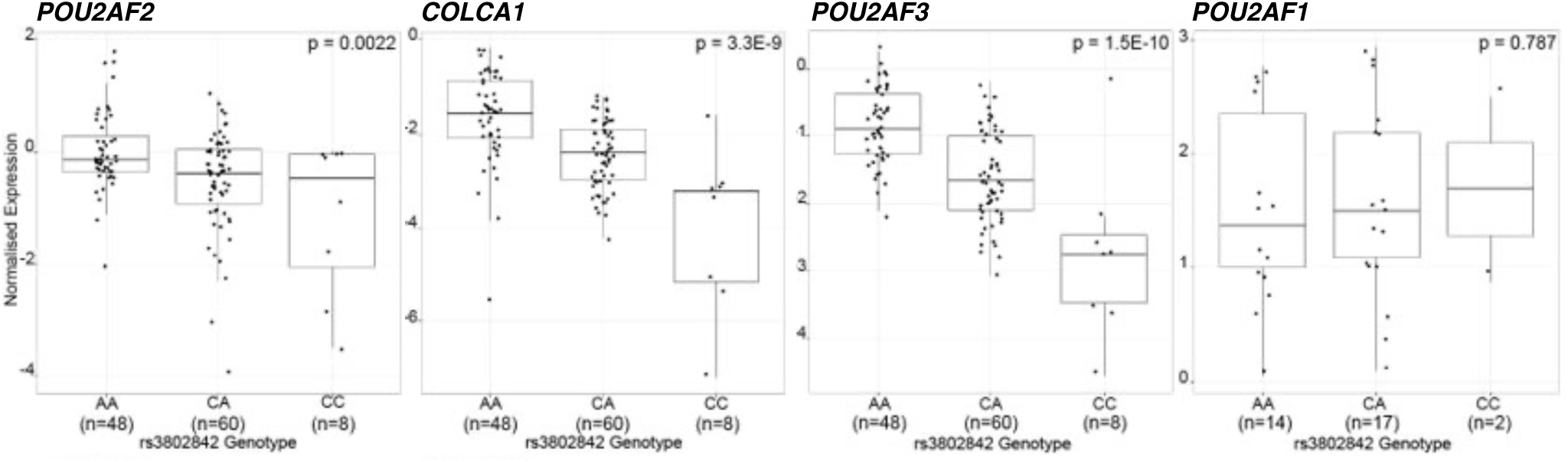
CRC risk-associated variation at 11q23.1 is correlated with the expression of *POU2AF2, COLCA1* and *POU2AF3* in the colorectal mucosa. qRT-PCR of 11q23.1 cis-eQTL targets *POU2AF2, COLCA1, POU2AF3* and nearby gene *POU2AF1,* in colorectal normal mucosa across tag variant rs3802842 genotype. C=risk, A=non-risk. P-values calculated by Spearman correlation rank sum test.

**Supplementary Figure S2.**
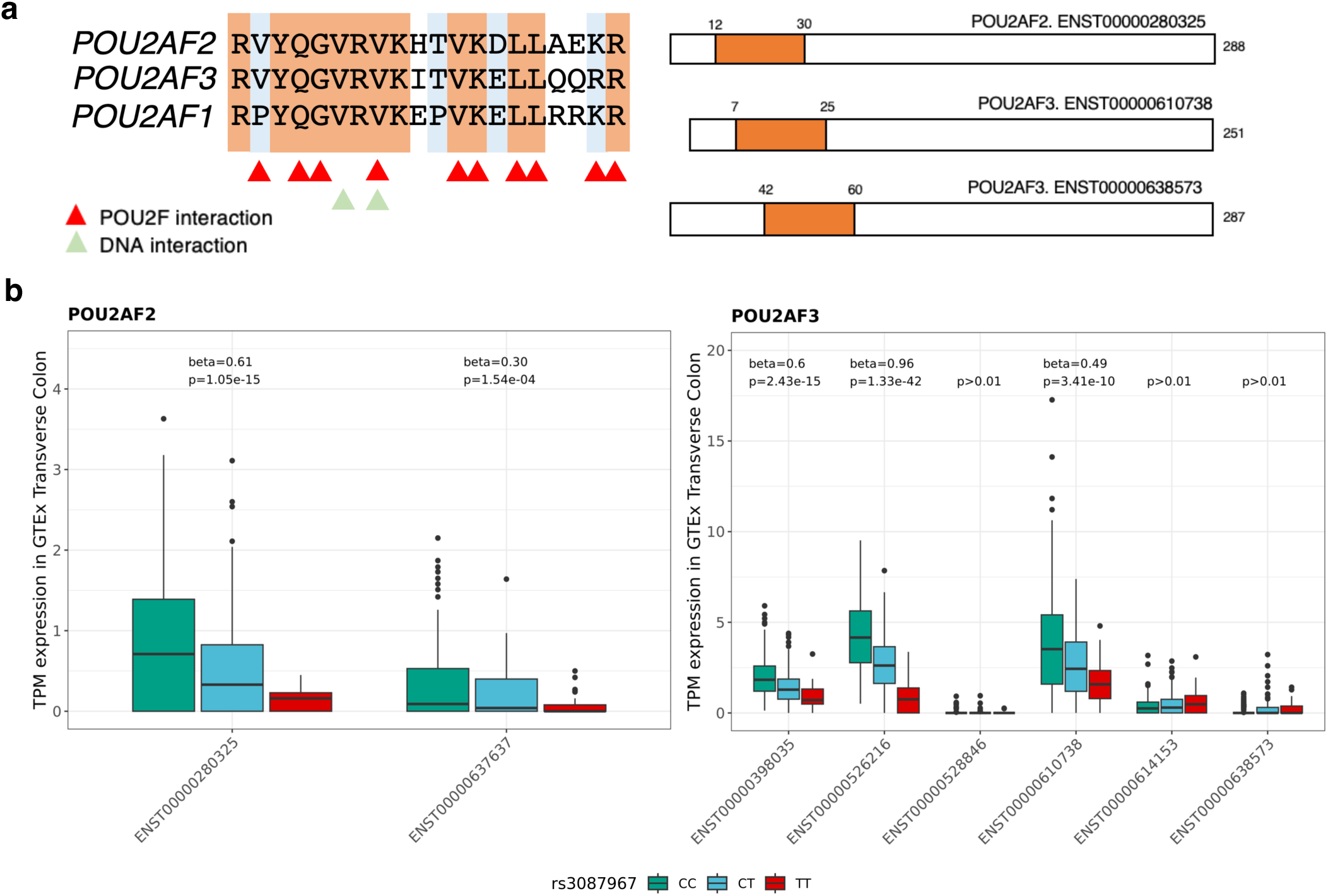
Transcript-specific expression and domain composition of *POU2AF2* and *POU2AF3.* (a) Sequence and encoding of POU2F interaction domains across *POU2AF2* and *POU2AF3* transcripts. Adapted from Wu *et al.,*^19^. (b) Expression of *POU2AF2* and *POU2AF3* transcripts across rs3087967 genotype in GTEx transverse colon. CC=174, CT=160, TT=33. Beta values and False discovery Rate (FDR) calculated by transcript-wide eQTL analysis. Orange highlighted transcripts encode POU2F interaction domain.

**Supplementary Figure S3.**
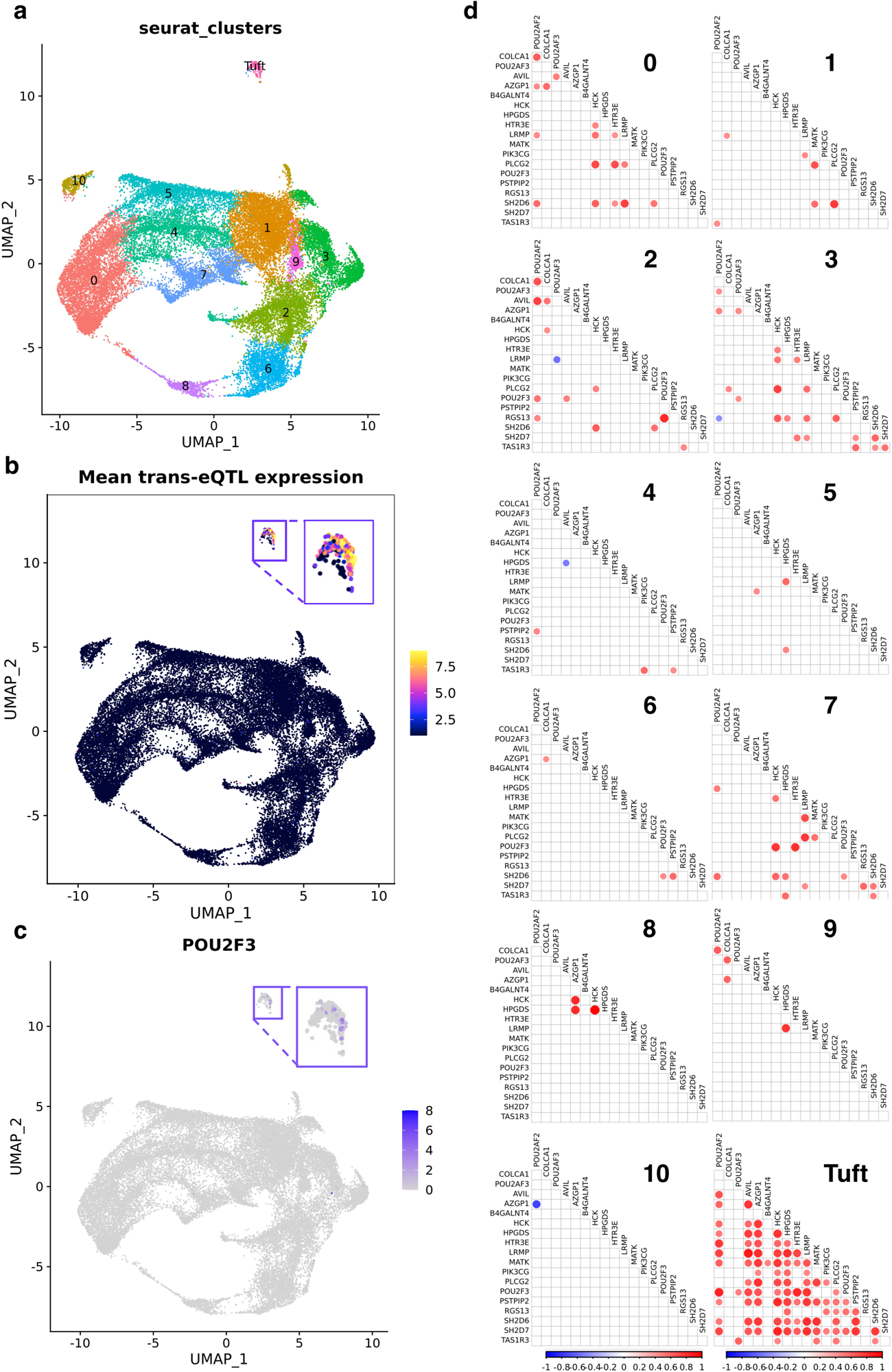
*POU2F3* exhibits tuft cell-specific correlation with *POU2AF2* in the human colon. (a) UMAP embedding of our previous analysis of Smillie *et al.,*^19^ healthy human colonic epithelium scRNAseq^14^. (b) The mean expression of refined trans-eQTL targets within cells. (c) *POU2F3* expression across cells. (d) Pairwise correlations between the expression of refined 11q23.1 trans-eQTL targets across scRNAseq clusters of (a). Only significant (p<0.05) associations are shown.

**Supplementary Figure S4.**
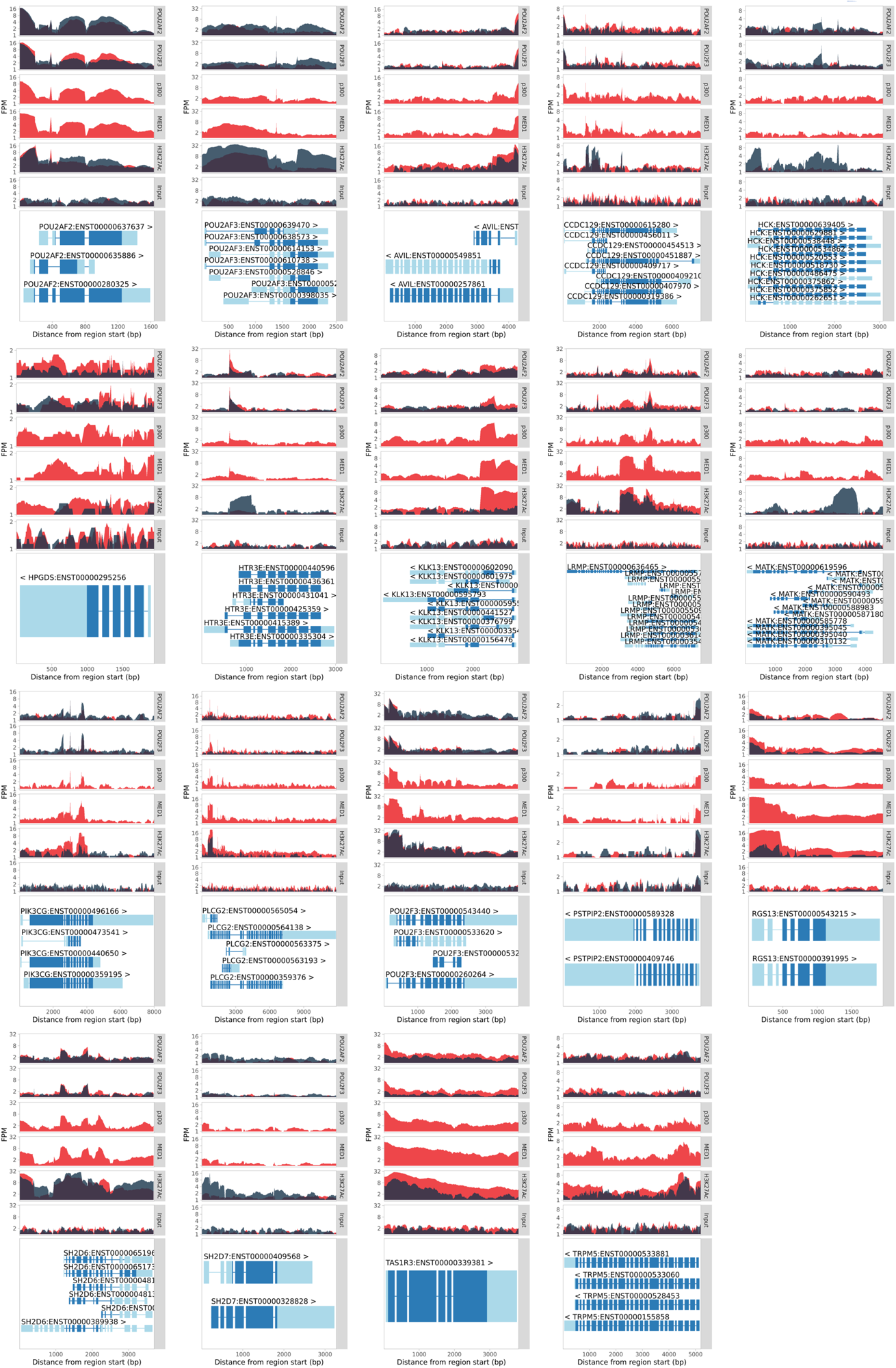
POU2AF2 and POU2F3 exhibit correlated binding at 11q23.1 trans-eQTL targets in SCLC cell lines indicative of transcriptional activation. POU2AF2 and POU2F3 enriched reads at *POU2AF2, POU2AF3* and 11q23.1 trans-eQTL targets bound by core motif sequence in Figure 4c. Individual replicates for each POU2AF2 antibody are merged within cell-line. Red histogram = NCIH211, Grey histogram = NCIH526.

**Supplementary Figure S5.**
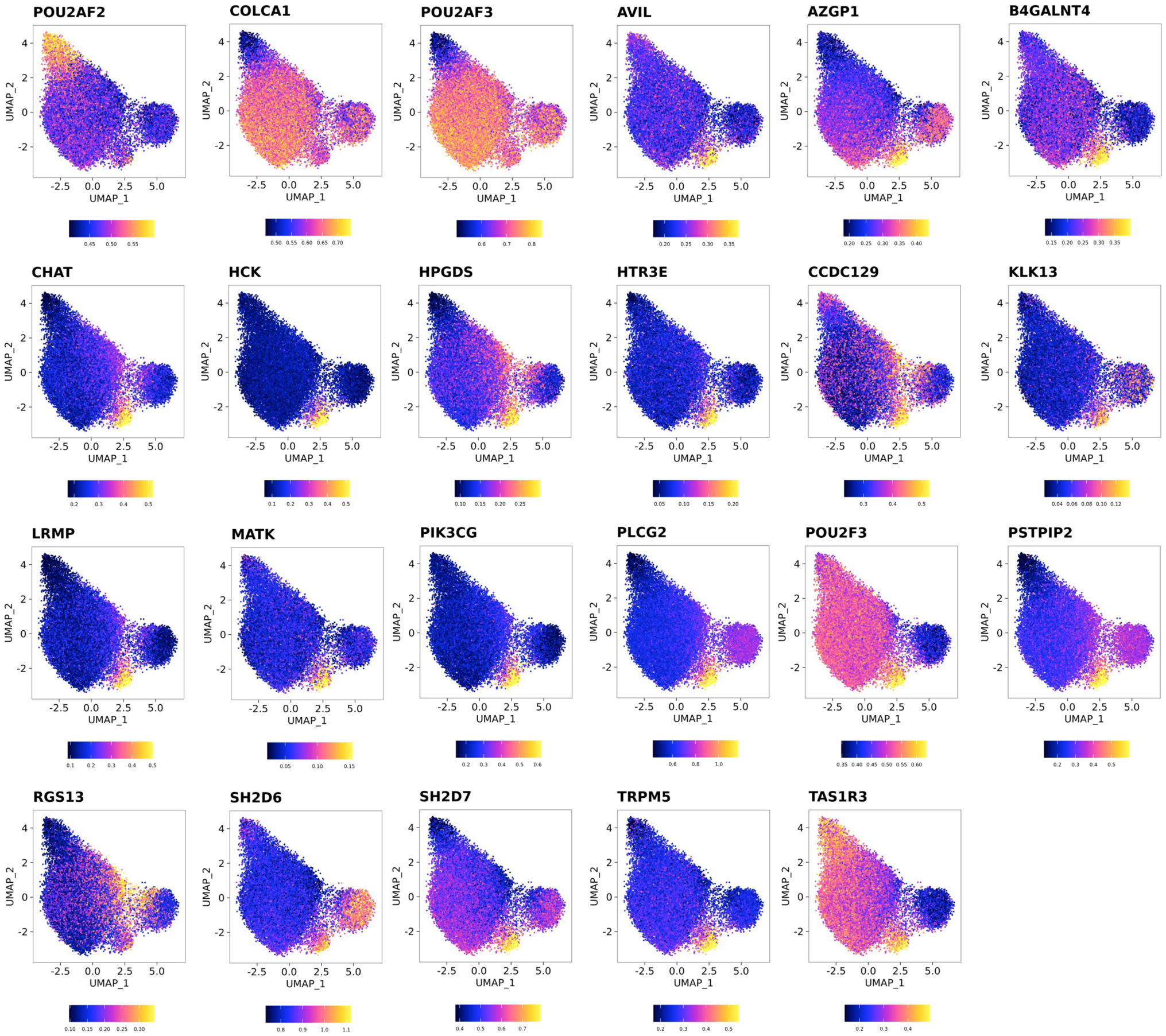
Chromatin at 11q23.1 trans-eQTL targets is exclusively accessible in tuft cells. Normalised accessibility of 11q23.1 cis- and refined trans-eQTL targets across individual cells.

**Supplementary Figure S6.**
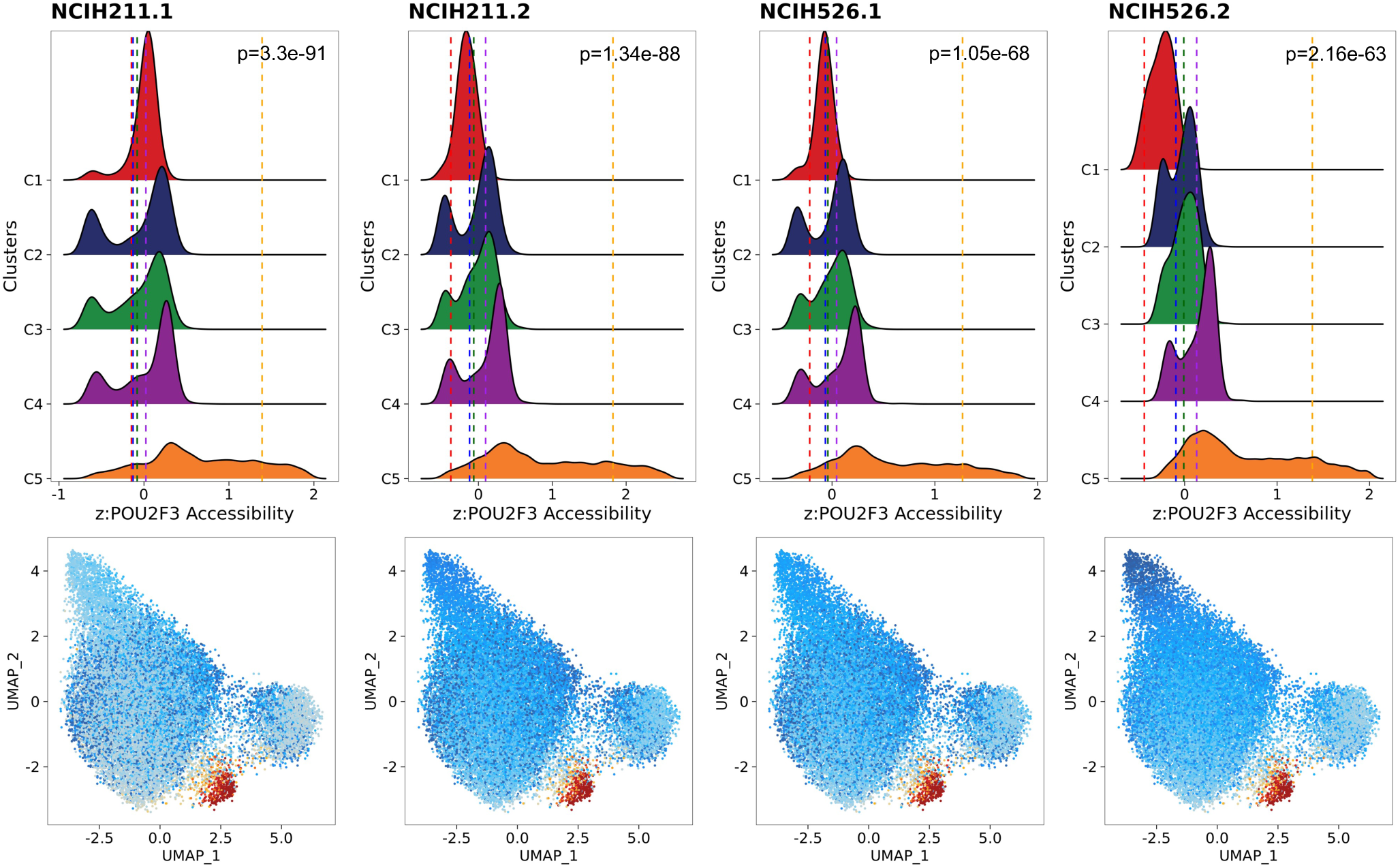
POU2F3-bound sequence accessibility is greatest in tuft-like cells. Relative accessibility of POU2F3-bound sequence accessibility across entire scATAC clusters (above) and individual cells (below). P-values calculated by t-test of normalised enrichment scores in cluster 5 compared to all other clusters combined.

**Supplementary Figure S7.**
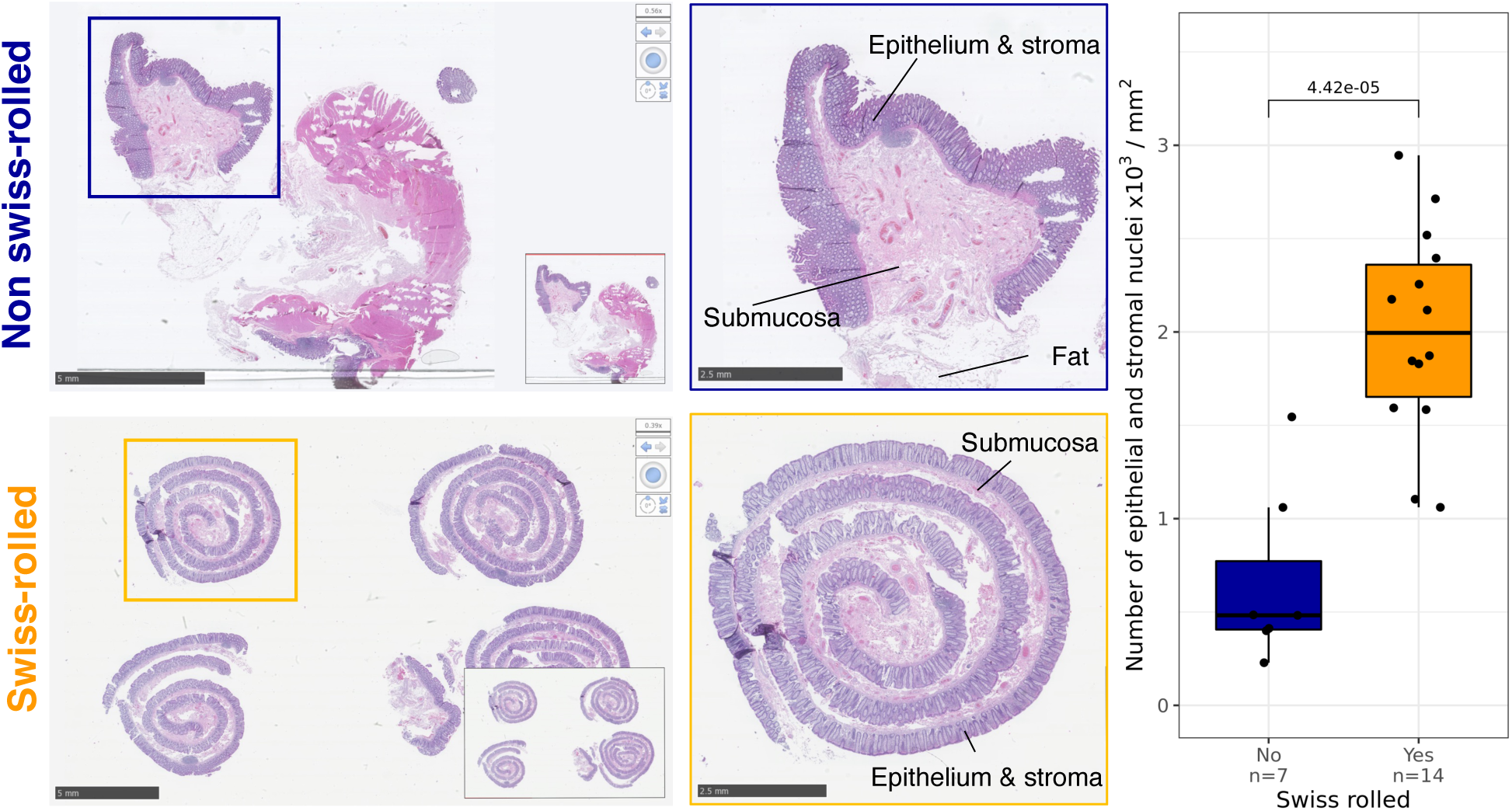
Optimisation of epithelial cell collection for detection of rare cell types. Haematoxylin and eosin stained samples of healthy colonic epithelium pre- (top) and post- (bottom) optimisation of ‘swiss-rolling’ method. Images taken using NDP View 2. P-value calculated by unpaired t-test.

**Supplementary Figure S8.**
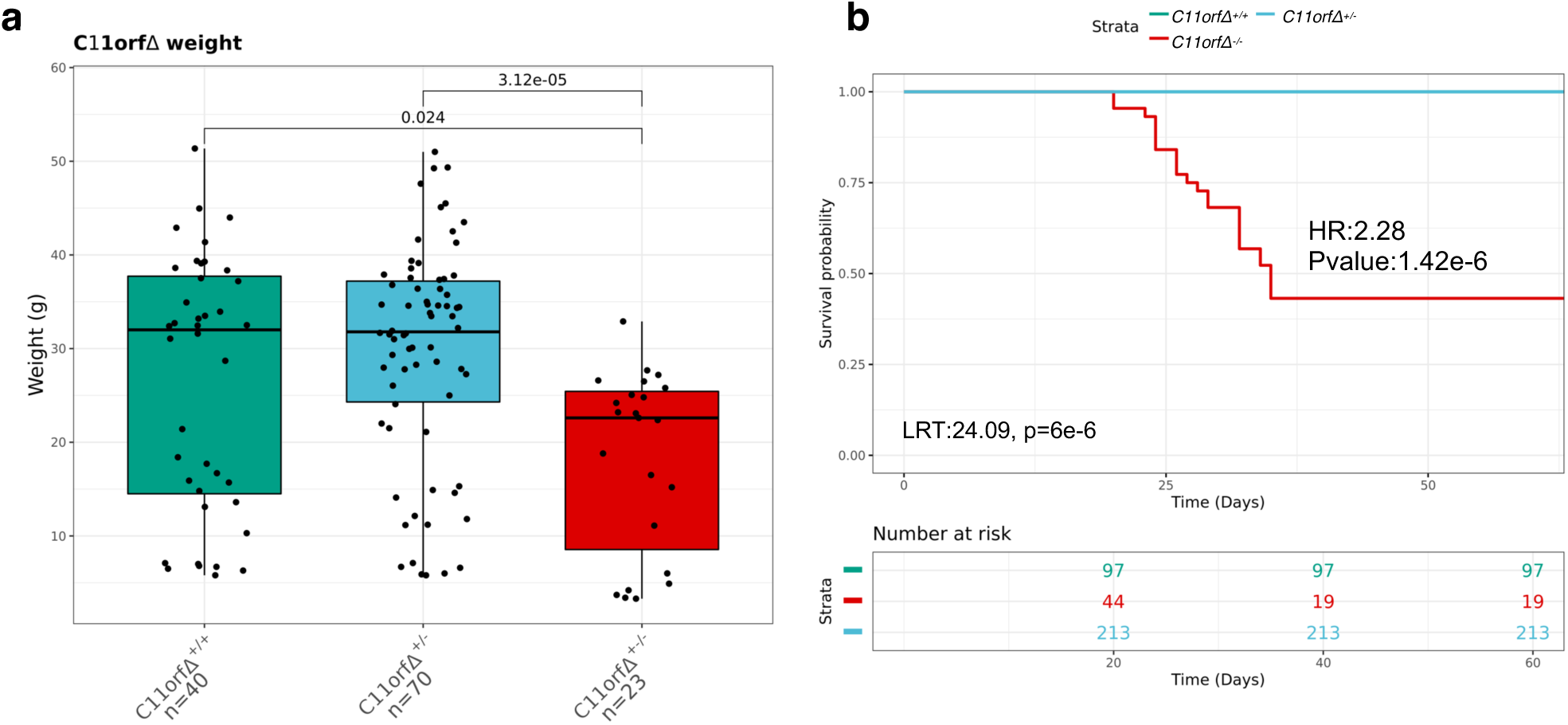
*C11orfΔ^-/-^* mice exhibit reduced weight and overall survival. Weight (a) and survival (b) of *C11orfΔ* mice. P values for weight comparisons are calculated by Benjamini-Hochberg correction of unpaired Wilcoxon ranks sum tests. Survival P-value calculated by cox proportional hazards model. LRT=Likelihood ration test, HR=hazard Ratio.

**Supplementary Figure S9.**
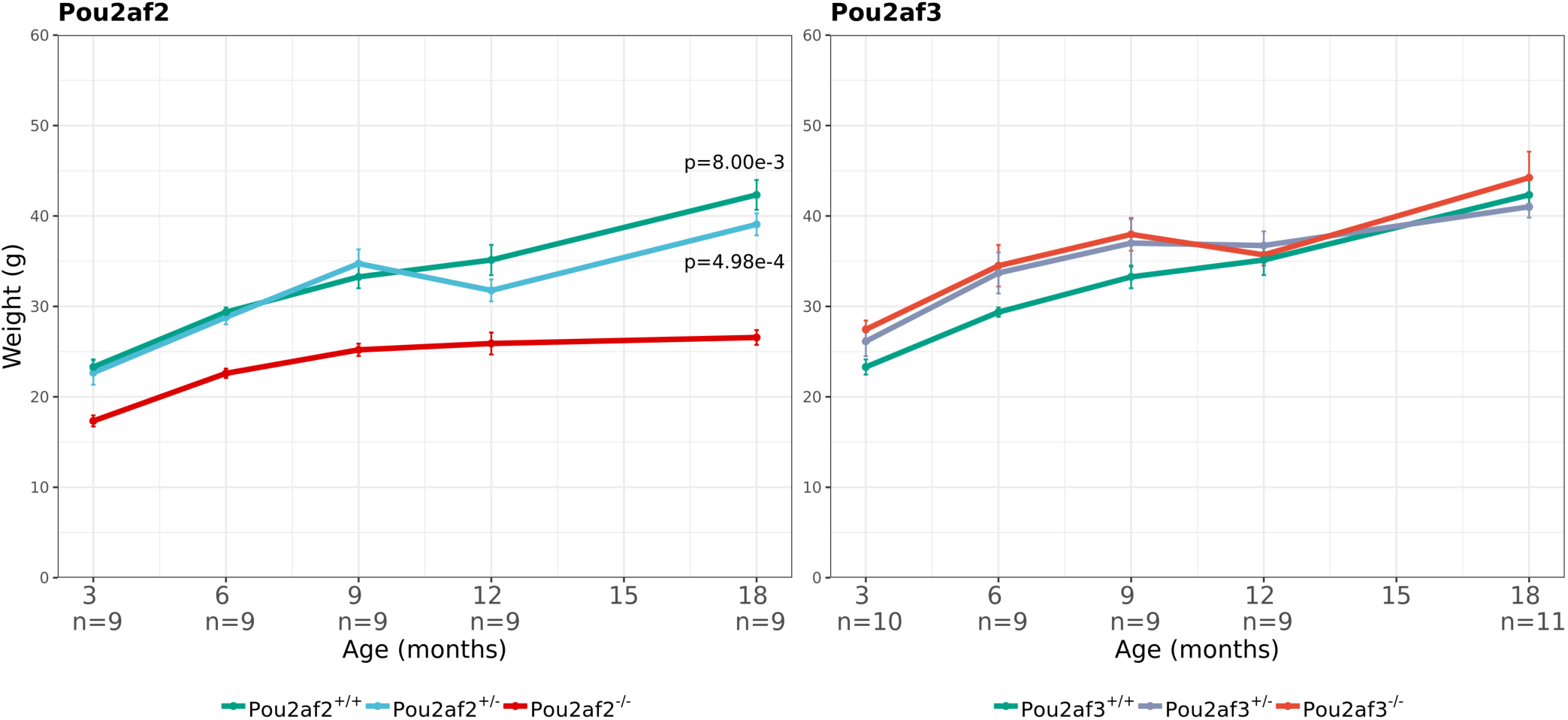
*Pou2af2^-/-^* but not *Pou2af3^-/-^* mice exhibit reduced weight. Error bars represent standard error about the mean. P-values are calculated by t-test of area under curve statistics for genotype comparisons against *Pou2af2^-/-^. Pou2af3* genotype was not associated with any change in weight.

